# Sleep-dependent clearance of brain lipids by peripheral blood cells

**DOI:** 10.1101/2025.10.30.685668

**Authors:** Bumsik Cho, Diane E. Youngstrom, Samantha Killiany, Camilo Guevara, Caitlin E. Randolph, Connor H. Beveridge, Pooja Saklani, Gaurav Chopra, Amita Sehgal

## Abstract

Sleep is typically viewed through a brain-centric lens, with little known about the role of the periphery. Here, we identify a sleep function for peripheral macrophage-like cells (hemocytes) in the *Drosophila* circulation, showing that hemocytes track to the brain during sleep and take up lipids accumulated in cortex glia due to wake-associated oxidative damage. Through a screen of phagocytic receptors expressed in hemocytes, we discovered that knockdown of *eater*, a member of the Nimrod receptor family, reduces sleep. Loss of *eater* also disrupts hemocyte adhesion to the brain and lipid uptake, which results in increased brain levels of Acetyl CoA and acetylated proteins, including mitochondrial proteins PGC1α and DRP1. Dysregulation of mitochondria, reflected in high oxidation and reduced NAD+, is accompanied by impaired memory and lifespan. Thus, peripheral blood cells, which we suggest are precursors of mammalian microglia, perform a daily function of sleep to maintain brain function and fitness.

## Main text

Sleep is a behavioral state shared by almost all animals. It is defined as a quiescent state associated with reduced consciousness that is different from coma or anesthesia because it is rapidly reversible with a stimulus^1,2^. The circadian system regulates sleep on a 24-hour cycle, but sleep is also regulated by homeostatic mechanisms, whereby the pressure to sleep increases with extended periods of wakefulness^1^. The importance of sleep is widely recognized, but the underlying mechanisms and functions are still debated.

While studies of sleep focus on the brain, sleep loss also has impact on the periphery^3^. In addition, there is now reason to believe that peripheral tissues can affect sleep^4,5^. The immune system is implicated in the control of sleep, particularly during sickness. In *Drosophila,* the NF-κB protein Relish acts in the fly fat body (functional equivalent of the liver) to regulate sleep following infection^6^. Sleep, in turn, influences recovery from bacterial and viral infections in both mammals and flies^7,8^. Sleep deprivation also increases the expression of sleep-promoting cytokines, such as TNFα or IL-6 and it may do so in the same way as inflammation, by increasing levels of the glucocorticoid hormone through the hypothalamic– pituitary–adrenal (HPA) axis or noradrenaline through the sympathetic nervous system (SNS)^7^. Thus, interactions between sleep and the immune system have been studied largely in pathological contexts, and have focused on signaling molecules. The role of circulating immune cells has not been addressed under pathological or normal physiological conditions. Importantly, peripheral mechanisms have never been implicated in the function of sleep.

Using *Drosophila* as a model system, we addressed a sleep function for circulating blood cells called hemocytes, 95% of which are macrophage-like plasmatocytes^9^ that function in immune responses. We show that at times of high sleep hemocytes localize to the brain and take up lipids accumulated in cortex glia. As lipid droplets in cortex glia reflect the transfer of wake-associated oxidative damage from neurons^10^, this uptake by hemocytes is expected to ease metabolic stress in the brain. Indeed, loss of the eater receptor, which mediates lipid uptake by hemocytes, causes increased acetylation in the brain, along with mitochondrial oxidation and reduced NAD+ levels. Thus, hemocytes, and eater in particular, act in a sleep-dependent fashion to maintain metabolic homeostasis in the brain.

### Hemocytes physically interact with the blood brain barrier at times of high sleep

To explore a possible interaction between hemocytes and the brain, we first utilized a tissue clearing method^11^ to visualize circulating hemocytes within the fly head (Fig. 1a, Extended Data Fig. 1a). We used the *HmlΔ-LexA* fly line to label hemocytes, and detected their localization throughout the head area, including the proboscis (pb), maxillary palp (mp), and ocellar (oc) regions, but not in the eyes or antenna (at) (Fig. 1a). To determine whether these populations of hemocytes mostly circulate or actually contact the brain, we dissected fly brains and visualized hemocytes using various markers (Fig. 1b-c, Extended Data Fig. 1b). Consistent with previous studies, Hml+ hemocytes were mostly located near the posterior part of the brain^12,13^, especially in the dorsally-located Pars Intercerebralis (PI) region. These Hml+ cells were also positive for other hemocyte markers like Srp-Hemo^14^ (Fig. 1b) and NimC1^15^ (Fig. 1c). Moreover, they were observed with other hemocyte Gal4 drivers, such as *Srp-Gal4*^16^, *Ppn-Gal4*^17^ and *Srp-Hemo-split-Gal4*^18^, and they expressed *eater-dsRed* that is specific to hemocytes^19^ (Extended Data Fig. 1b). Hemocytes were not detected near the ventral nerve cord or inside the brain (Extended Data Fig. 1c). Based on these observations, we conclude that hemocytes circulate in the fly head cavity and possibly interact with the brain at specific sites.

**Figure 1.**
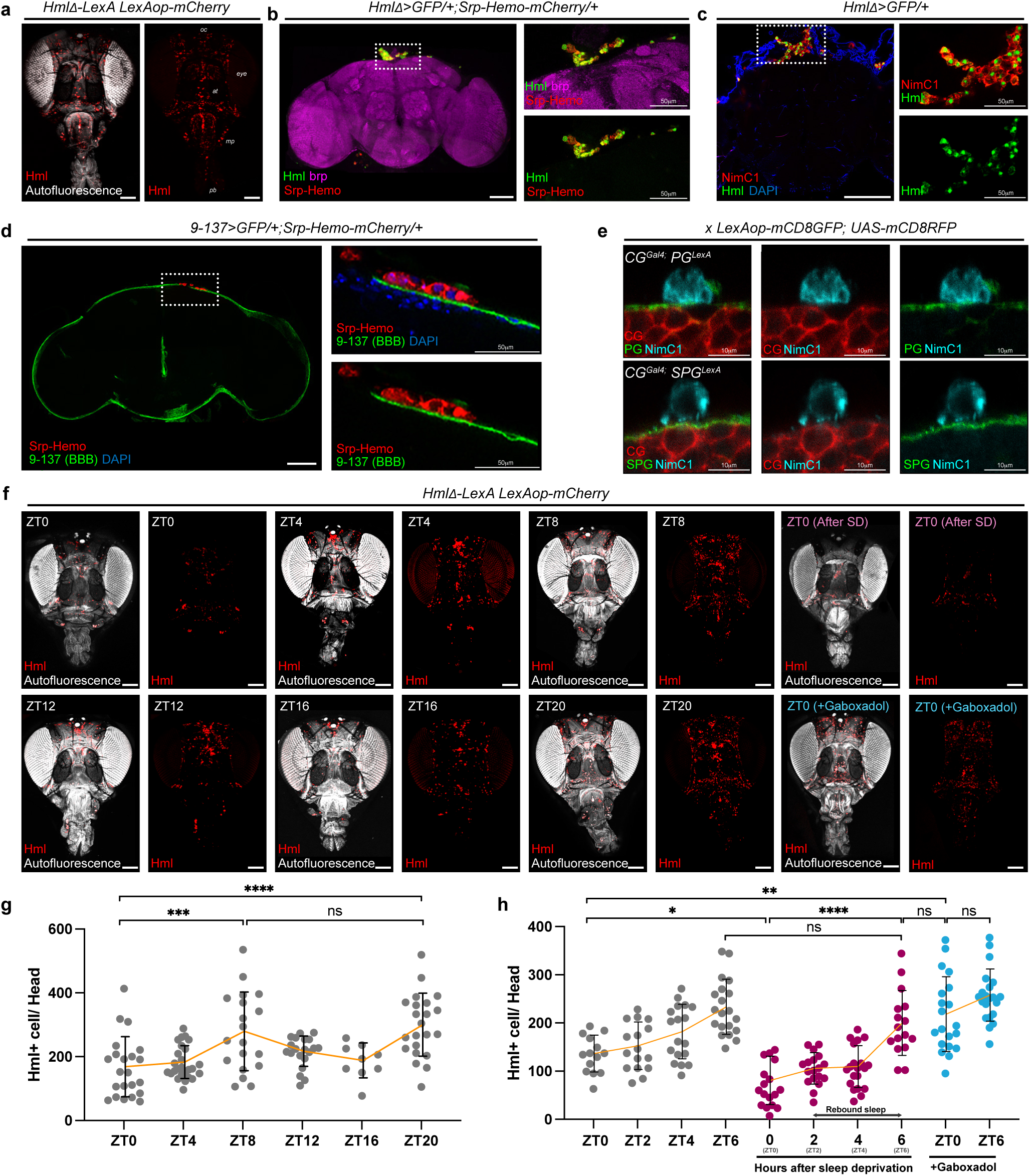
Blood cells (hemocytes) circulate in the fly head cavity and adhere to the BBB. **a.** Localization of Hml-LexA+ hemocytes (*Hml Δ-LexA LexAop-mCherry*, red) within the fly head cavity. The structure of the head was visualized using a UV laser (autofluorescence, white). Labels indicate the ocelli (oc), antenna (at), maxillary palp (mp), and proboscis (pb). **b-c.** Location of hemocytes near the brain. Hml+ (green) or Srp-hemo+ (red) hemocytes are located on the top middle area of the brain (left). Magnified images on the right. Dotted box indicates magnified area. Structure of the brain is visualized with brp (magenta) **(b)**. Hml+ (green) or NimC1+ (red) hemocytes are seen in the same area as (b) (left). Magnified images on the right. Dotted box means magnified area **(c)**. **d-e.** Localization of hemocytes near the blood brain barrier. Srp-hemo+ (red) hemocytes are located next to the BBB (*9-137-Gal4 UAS-GFP*, green). Magnified images on the right. Dotted box means magnified area **(d)**. NimC1+ (cyan) hemocytes are located next to the perineurial glial cell (PG) (*R85G01-LexA LexAop-mCD8GFP,* green) (top) or sub-perineurial glial cell (SPG) (*R54C07-LexA LexAop-mCD8GFP,* green) (bottom). Cortex glia (CG) is visualized with RFP (*NP2222-Gal4 UAS-mCD8RFP,* Red) **(e)**. **f-h.** Visualizing of circulating hemocytes in the head at different times of day. *Hml-LexA*+ hemocytes (*Hml Δ-LexA LexAop-mCherry*, red) are visualized at Zeitgeber Time (ZT) times ZT 0, 4, 8, 12, 16, and 20. Also from ZT 0 to 6 following 12 hours of sleep deprivation or at ZT 0 and 6 following 2mM Gaboxadol feeding. Compound image showing autofluorescence (white) with hemocytes (red) is on the left side and hemocytes only (red) on the right **(f)**. Count of *Hml-LexA*+ hemocytes per head at different times of day (data shown in F) **(g)**. Count of *Hml-LexA*+ hemocytes per head after 12 hours sleep deprivation or 2mM Gaboxadol feeding (data shown in f) **(h)**. Yellow line indicates average of the number of hemocytes. Tukey’s multiple comparisons test was performed for data analysis. DAPI: blue, White scale bar, 100 μm unless otherwise indicated. ns: not significant (p>0.01); *p<0.1; **p<0.01, ***p<0.001. ****p<0.0001. Bars in graphs: the median with standard deviation.

The fly brain is separated from the periphery by the blood-brain barrier (BBB)^20^. To assess whether hemocytes near the PI region interact with the BBB, we visualized hemocytes together with the BBB-specific Gal4 line that marks both perineurial glia (PG) and sub-perineurial glia (SPG) cells of the BBB^21^ (Fig. 1d, Extended Data Fig. 1b), or using Gal4 lines that individually mark PG cells (*NP6293-Gal4*) or SPG cells (*moody-Gal4*) (Fig. 1e, Extended Data Fig. 1d-e). We found that Srp-Hemo+, eater+, or NimC1+ hemocytes were located adjacent to the BBB (Fig. 1d-e, Extended Data Fig. 1b, d-e). Moreover, when we visualized hemocytes together with PG markers, we observed that extensions of the PG membrane physically contact hemocytes (Fig. 1e, Extended Data Fig. 1f). Although SPG and hemocytes are separated by PG cells, hemocyte membranes are also in contact with SPG cells (Fig. 1e). Indeed, we used the GFP Reconstitution Across Synaptic Partners (GRASP) technique^22^ to confirm direct physical interaction between hemocytes and SPG cells (Extended Data Fig. 1g, h). Similar direct interaction of SPG cells with hemocytes was observed in pupal stages via electron microscopy (EM)^23^. A previous study found that cortex glia (CG) cells contact or share membranes with SPG^24^ cells, so it is possible that hemocytes also directly contact CG cells but we did not detect a GRASP signal (data not shown). Based on these findings, we conclude that hemocytes exist within the fly head cavity and physically interact with glial cells, particularly glia of the BBB.

We also asked if hemocyte recruitment to the brain is influenced by the sleep:wake cycle (Fig. 1f-g). At Zeitgeber Time (ZT)8 and ZT20 (ZT0=lights on, in circadian terms), which are times of the afternoon siesta and night-time sleep respectively^25^, the number of hemocytes in the head was higher than at other times of day (Fig. 1f-g). To confirm sleep-dependent hemocyte recruitment to the fly head, we compared hemocyte numbers following sleep deprivation or gaboxadol feeding to induce sleep (Fig. 1f, h). Sleep deprivation reduced hemocyte numbers in the head, but the numbers recovered during rebound sleep (Fig. 1f, h). In contrast, feeding gaboxadol increased hemocyte numbers in the fly head, with no significant differences across ZT time points (Fig. 1f, h). From these results, we surmise that the interaction between hemocytes and the brain is increased during sleep.

### eater expressed in hemocytes is required for sleep regulation

Given that hemocytes circulate in the fly head and are more abundant during sleep than wake, we hypothesized that the function of hemocytes may be relevant for sleep. To use an unbiased approach towards such function, we examined recent single-cell RNA sequencing data^26^ for transcripts expressed highly in hemocytes. Analysis of the biological functions of the top 100 genes in hemocytes through g:Profiler^27^ revealed significant annotations for defense response against other organisms, phagocytosis, and immune system processes. Since the phagocytosis of gram-positive or negative bacteria in *Drosophila* is typically mediated by Nimrod receptor family genes^15^, we focused on a possible role for this family (*NimA*, *NimB1-5*, *NimC1-4*, *drpr*, *eater*, *Col4a1*, *PGRP-LC*) in the regulation of sleep. In addition, because we observed that more hemocytes are localized in the head cavity during the sleep state in flies (Fig. 1f-h), we also added genes previously identified to be involved in hemocyte migration. These include activin-β signaling factors (*babo*, *put*, *Smox*) that are important for sessile localization of hemocytes^28^, or the PDGF/VEGF signaling receptor (Pvr) that functions in embryonic blood cell migration^29^. Lastly, we included genes associated with lipid uptake (*Lsd-1, Lsd-2, GLaz, Karl, Nplp2, Apoltp, Acsl, LpR1, LpR2, eater, crq, apolpp*), as previous studies have shown that hemocyte functions are linked to lipid uptake^30,31^, processing^32,33^, and clearance^34,35^, which are critical for immune system activation, animal growth, and metabolism.

We then knocked down each of the genes above using RNAi and assayed effects on sleep. Knockdown of the gene *eater* reduced sleep, validated using two different *eater* RNAi lines (Extended Data Fig. 2a) with two distinct hemocyte Gal4 drivers (Extended Data Fig. 2b-c). To further confirm the *eater* knockdown phenotype, we assessed the sleep patterns of *eater* null mutants. Both male and female *eater* mutant flies exhibited reduced daytime and nighttime sleep (Fig. 2a-b), with shorter sleep bout lengths and more sleep bouts during the nighttime but no reduction in activity counts during wake (Fig. 2b). This phenotype indicates that the reduced sleep of the *eater* mutants is also fragmented and not associated with motor activity impairment. Reduction of sleep was not observed in *eater* heterozygous mutants (Extended Data Fig. 2d), indicating that the effect is recessive. And transgenic expression of *eater* in hemocytes rescued the mutant sleep phenotype, confirming that the phenotype maps to the *eater* gene (Fig. 2c). We attempted to also rescue *eater* with a homologous mammalian protein, MEGF11^36^, but this was not successful (Extended Data Fig. 2e), perhaps because MEGF11 contains only 17 EGF-like repeats, while *eater* has 32.

**Figure 2.**
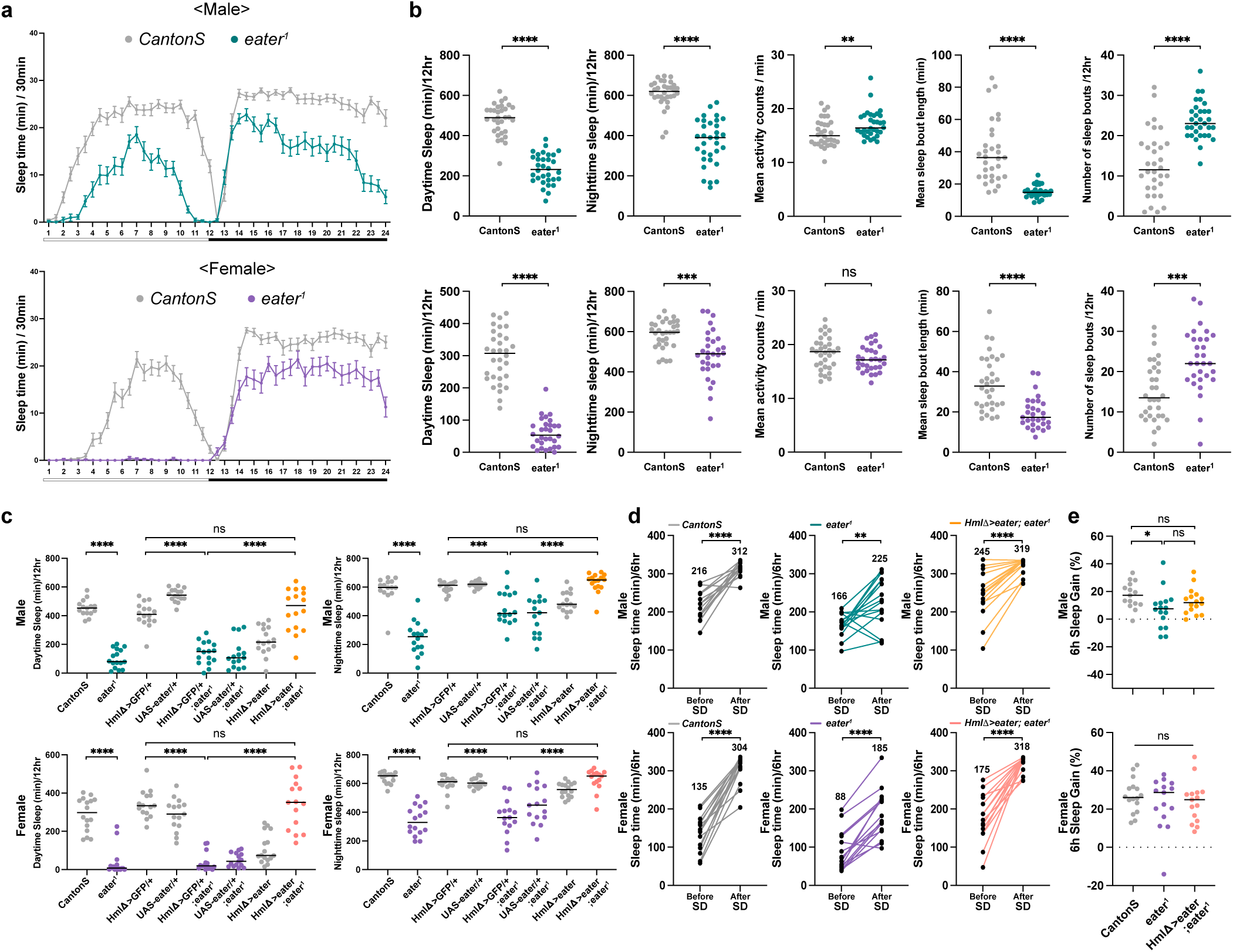
eater in hemocytes regulates sleep. **a-b.** Sleep analysis of wild-type (*CantonS*) and *eater* mutant (*eater*^1^) flies. Graphs indicating sleep amount in wild-type and *eater* mutant flies for both males (top, dark green) and females (bottom, purple) **(a)**. Quantification of daytime sleep, nighttime sleep, mean activity counts per minute, mean sleep bout length, and number of nighttime sleep bouts in males (top, dark green) and females (bottom, purple) **(b)**. Mann-Whitney test was performed for data analysis. **c-e.** Rescue of sleep reduction in *eater* mutants through hemocyte specific *eater* expression. Quantification of total sleep time, showing *eater* mutants (dark green dots for males, purple dots for females) and *eater* overexpression in mutants (yellow dots for males, pink dots for females) **(c)**. Comparison of time of sleep during and after sleep deprivation in males (top) and females (bottom). The numbers above the graph represent the total sleep time for each genotype for 6 hours **(d)**. Analysis of rebound sleep at ZT 0-6 before and after sleep deprivation (gain in sleep over this time), with data for males (top, dark green) and females (bottom, purple) **(e)**. Tukey’s multiple comparisons test was performed for data analysis in (c), (e). Paired t test was performed for data analysis in (d). ns: not significant (p>0.01); *p<0.1; **p<0.01, ***p<0.001. ****p<0.0001. Bars in graphs: the median with SEM in (a). White and black bar below the graph in (a) represents day (white) and night (black).

To exclude the possibility that the reduced sleep phenotype comes from developmental effects, we used a temperature-sensitive Gal80 (*Hml∆-Gal4 UAS-GFP, Tub-Gal80ts*) to block Gal4 expression during development. Knocking down *eater* only in the adult stage was enough to decrease sleep indicating that the phenotype is not due to developmental effects (Extended Data Fig. 2f). Furthermore, lack of a circadian phenotype under constant dark conditions (Extended Data Fig. 2g) demonstrated that the decreased sleep amount in *eater* mutants is not driven by circadian rhythm alterations, suggesting that *eater* more directly affects sleep.

To assess whether *eater* mutants exhibit rebound sleep, flies were subjected to sleep deprivation, and the amount of sleep gained during recovery was compared to that in control flies (Fig. 2d-e). While the total sleep during a 6-hour recovery period was lower in *eater* mutants than in controls, the percentage of sleep gain was comparable across groups (Fig. 2d-e). These results indicate that homeostatic regulation of sleep is intact in *eater* mutants. Altogether, we conclude that *eater* in hemocytes is required to maintain daily sleep in adult flies.

### Hemocyte transfer rescues the eater mutant sleep phenotype

Although *eater* is known to be expressed specifically in hemocytes^37^, we further tested whether the mutant sleep phenotype derives from hemocytes by transferring wild-type hemocytes to *eater* mutant flies to determine whether this could restore their normal sleep pattern. Due to the challenge of obtaining pure hemocytes from adult flies without any enzymatic treatment, we used larval hemocytes for the transfer experiment (Extended Data Fig. 3a). Using the tissue clearing method, we confirmed that transferred labeled hemocytes were properly circulating throughout the body and head regions of wild-type and *eater* mutant flies 4h after injection (Extended Data Fig. 3b-c).

We then assessed whether the hemocyte transfer rescued the sleep phenotype in *eater* mutants. We injected hemocytes at the beginning of the day, ZT2, to minimize wound-induced increases in sleep that occur with injection at night^6^. In wild-type flies, transfer of either wild-type (WT) or *eater* mutant hemocytes did not alter sleep amount or patterns (Fig. 3a, c). In *eater* mutants, injection alone (e.g. with PBS) elicited a small, although insignificant, increase in sleep. However, injection with wild type hemocytes produced the most robust sleep increase that was significantly higher than in flies injected with PBS or *eater* mutant hemocytes (Fig. 3b-c). The fact that both WT hemocyte transfer to *eater* mutants (Fig. 3b-c) and genetic restoration of *eater* expression in the hemocytes of *eater* mutants (Fig. 2c) are able to rescue the sleep loss phenotype confirms that *eater* function in hemocytes is necessary and sufficient to regulate sleep.

**Figure 3.**
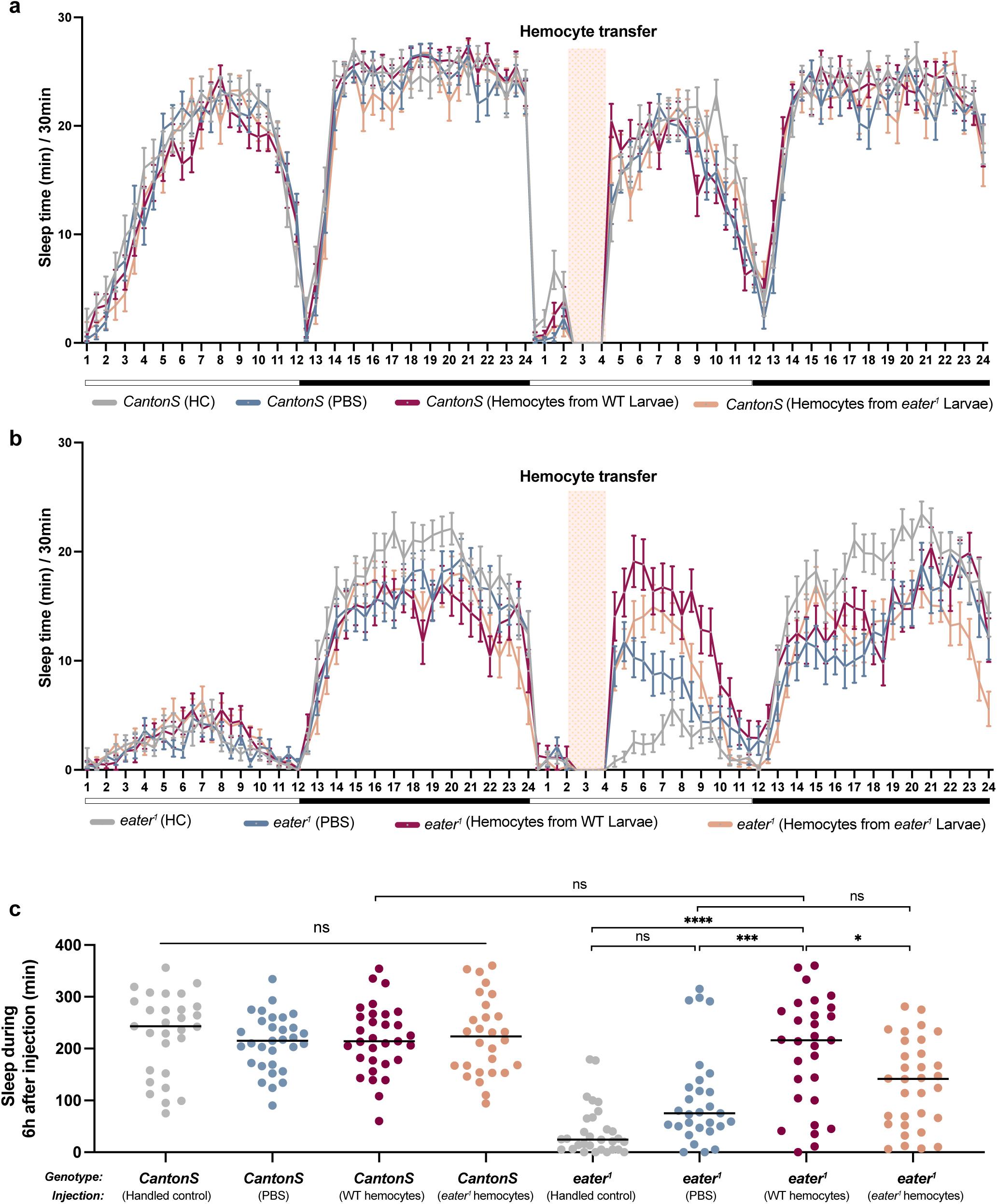
Hemocyte transfer rescues the eater mutant sleep phenotype. **a-c.** Comparison of sleep amount in wild-type (*CantonS*) and *eater* mutant (*eater*^1^) before and after larval hemocyte transfer. Profile of sleep in wild-type (*CantonS*) before and after hemocyte transfer **(a)**. Profile of sleep in the *eater* mutant (*eater*^1^) before and after hemocyte transfer **(b)**. Quantitative analysis of sleep measured during the 6 hours after hemocyte transfer in both wild-type and *eater* mutant flies **(c)**. HC: hand control (grey; not wounded but exposed to CO_2_), PBS: PBS injection (blue), WT (dark red; *CantonS* hemocyte transfer), *eater*^1^ (orange; *eater*^1^ hemocyte transfer). Tukey’s multiple comparisons test was performed for data analysis. ns: not significant (p>0.01); *p<0.1; **p<0.01, ***p<0.001. ****p<0.0001. Bars in graphs: the median with SEM. White and black bar below the graph in (a) and (b) represents day (white) and night (black). The red shade in (a) and (b) indicates the time point when hemocytes were transferred.

### Eater protein in hemocytes clears lipids from cortex glia

The Eater protein, which contains 32 EGF-like repeats, is known to be involved in three key functions: 1) Phagocytosis of Gram-positive bacteria^38^, 2) Cell-to-cell adhesion^39^ , and 3) Low density lipoprotein (LDL) uptake^37^. These functions are also conserved in mammalian proteins containing EGF-like repeats^40^. Because we did not deliver bacterial challenges to the fly, we investigated the other two functions of Eater.

First, we assayed the number of circulating hemocytes in the fly head cavity in the *eater* knockdown background and found that the number was reduced relative to wild-type (Fig. 4a-b). The number of head hemocytes was rescued when sleep was increased by gaboxadol feeding but with high variability from fly to fly (Fig. 4a-b), which could reflect impaired adhesion to glial cells. Thus, we also examined whether the loss of *eater* in hemocytes affects their adhesion to glial cells at the brain surface (Extended Data Fig. 4a-c). Both in *eater* knockdown and *eater* mutants, fewer hemocytes were observed near glial cells, consistent with the reduced number in the head cavity (Extended Data Fig. 4a-c), and this was rescued by re-introducing *eater* expression in hemocytes (Extended Data Fig. 4c). Given that this manipulation was also sufficient to rescue sleep, together with the brain association of hemocytes at times that correspond to sleep (Fig. 1f-h, Fig. 4a-b, Extended Data Fig. 4a-c), we conclude that hemocytes recruitment and adhesion to glial cells influences sleep in flies.

**Figure 4.**
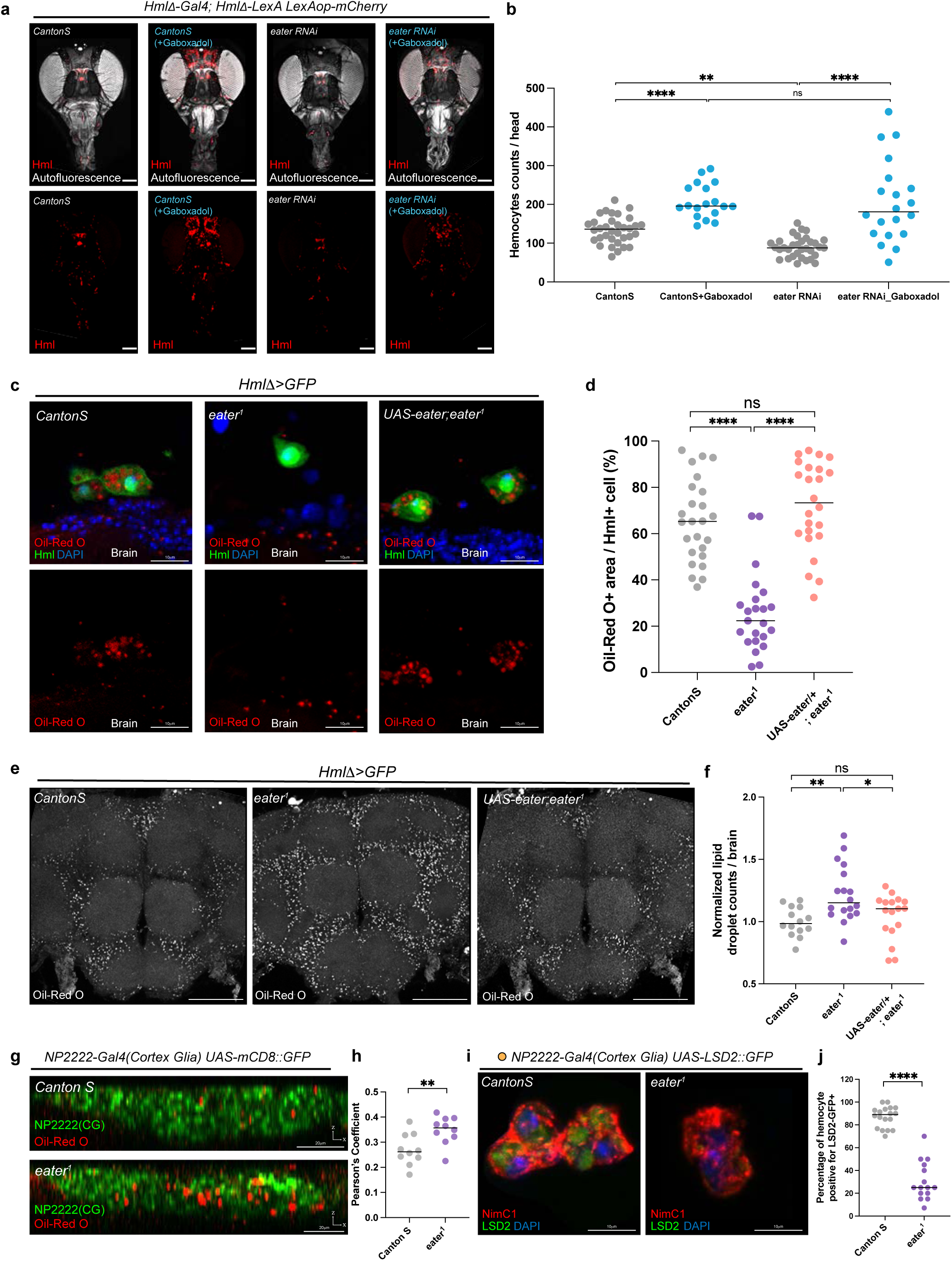
Eater is required for hemocytes to take up lipids from cortex glial cells. **a-b.** Comparison of circulating hemocytes in the head upon *eater* knockdown. Wild type or *eater* knockdown *Hml-LexA*+ hemocytes (*Hml Δ-Gal4; Hml Δ-LexA LexAop-mCherry*, red) are visualized at ZT2 or after 2mM Gaboxadol feeding **(a)**. Quantification of circulating hemocytes on **(b)**. Tukey’s multiple comparisons test was performed for data analysis. **c-d.** Visualization of lipid droplets in Hml+ (green) hemocytes stained with Oil-Red O (red) in wild-type (*HmlΔ>GFP/+*), *eater* mutant (*HmlΔ>GFP/+; eater*^1^), and *eater* rescue background (*HmlΔ>GFP, UAS-eater/+*; *eater*^1^) **(c)**. Quantification of Oil-Red O positive area within individual hemocytes **(d)**. Tukey’s multiple comparisons test was performed for data analysis. **e-h.** Visualization of lipid droplets in the brain stained with Oil-Red O (gray) in wild-type (*HmlΔ>GFP/+*), *eater* mutant (*HmlΔ>GFP/+; eater*^1^), and *eater* rescue background (*HmlΔ>GFP, UAS-eater/+; eater*^1^). *eater* mutants exhibit more lipid droplet accumulation **(e)**. Quantification of Oil-Red O positive lipid droplets in the brain (**f)**. Observation of Oil-Red O positive lipid droplets (red) in the brain with a cortex glia cell marker in wild-type (top, *NP2222-Gal4 UAS-mCD8GFP/+*) or *eater* mutant flies (bottom, *NP2222-Gal4 UAS-mCD8GFP/+; eater*^1^) **(g)**. Pearson’s coefficient of Oil-Red O co-localization with cortex glia marker **(h)**. Tukey’s multiple comparisons test was performed for data analysis in (f). Unpaired t test was performed for data analysis in (h). **i-j.** Observation of lipid droplets in hemocytes originated from the glia. LSD2-GFP positive lipid droplet from the cortex glia (green) were detected in the NimC1 positive (red) hemocytes. Hemocyte from the control (left, *NP2222-Gal4 UAS-LSD2::GFP/+*) contain more lipid droplets than those from the *eater* mutant (right, *NP2222-Gal4 UAS-LSD2::GFP/+; eater*^1^) **(i)**. Quantification of the LSD2::GFP+ area within individual hemocytes **(j)**. Mann-Whitney test was performed for data analysis. ns: not significant (p>0.01); *p<0.1; **p<0.01, ***p<0.001. ****p<0.0001. Bars in graphs: the median. DAPI: blue, White scale bar, 100 μm unless it’s not indicated.

Next, we investigated whether hemocytes take up lipid droplets through Eater. First, we visualized lipid droplets (LD) using a GFP-tagged LD domain^41^ in hemocytes (Extended Data Fig. 4d). We found that hemocytes were positive for Oil-Red O staining, and this staining co-localized with LD-GFP (Extended Data Fig. 4d). Additionally, we confirmed that lipids in the hemocytes were positive for BODIPY (Extended Data Fig. 4e). Interestingly, Oil-Red O staining in hemocytes was significantly reduced in *eater* mutants (Fig. 4c-d). This reduction in Oil-Red O staining in hemocytes was restored when Eater was rescued in hemocytes (Fig. 4c-d), indicating that *eater* is important for LD uptake into hemocytes.

We showed previously that LD accumulation in glial cells changes over the sleep:wake cycle and increases following sleep deprivation^10^. Given that *eater* mutants exhibit reduced sleep compared to wild-type flies and less lipid accumulation in hemocytes, we examined LD accumulation in brains (Fig. 4e-f). Compared with wild-type, Oil-Red O-positive LDs were increased in glial cells of *eater* mutants (Fig. 4e-f) or with *eater* knock-down in hemocytes (Extended Data Fig. 4f-g), and their accumulation was reduced when *eater* was rescued in hemocytes (Fig. 4e-f). Consist with the previous report^10^, we found that most lipid droplets accumulate in cortex glia, with lower levels in the BBB (Fig. 4g-h, Extended Data Fig. 4h-i). LD accumulation in the BBB also appeared to be higher in *eater* mutants, suggesting that these cells are also affected by loss of Eater.

We next tested the hypothesis that LDs in hemocytes are derived from glia. Thus, we expressed GFP-tagged Lipid Storage Droplet2 (LSD2) (*UAS-LSD2::GFP*)^42,43^ in glial cells using the pan-glial driver (*Repo-Gal4*) and checked whether LSD2::GFP was transferred to hemocytes. More than 80% of hemocytes displayed a glial cell derived LSD2::GFP signal, demonstrating that hemocytes take up lipid droplets from glial cells (Extended Data Fig. 4j-k). To determine which glial subpopulations transfer LDs to hemocytes, we used specific glial drivers. Given that LDs are known to accumulate in cortex glia, it was not surprising to find that approximately 75% of hemocytes were LSD2::GFP-positive when using a cortex glia driver (*NP2222-Gal4*), a level comparable to that observed with the pan-glial driver (Extended Data Fig. 4j-k). This suggests that cortex glia are the predominant glial cells transferring LDs to hemocytes. In contrast, when using drivers specific for other glial subpopulations, approximately 50% of hemocytes were LSD2::GFP-positive with an astrocyte-like glia driver (*Alrm-Gal4*), ∼20% with an ensheathing glia driver (*MZ0709-Gal4*), and ∼10% with a BBB glia driver (*9-137-Gal4*) (Extended Data Fig. 4j-k). When LSD2::GFP was expressed with the cortex glia driver (*NP2222-Gal4*) in the *eater* mutant background, fewer LSD2::GFP droplets were observed in hemocytes (Fig. 4i-j). Overall, we conclude that hemocytes interact with glial cells, particularly cortex glia, to uptake LDs through the Eater protein. If the cell-adhesion or LD uptake function of Eater is diminished, then excess lipids accumulate in cortex glia.

### Eater mediates the uptake of acetylated lipoproteins

Using a multiple reaction monitoring (MRM)-based screening lipidomic approach^44,45^, we profiled lipids in hemocytes from the head cavity (Extended Data Fig. 5a-c). We found that phosphatidylcholines (PCs) and phosphatidylethanolamines (PEs) exhibit the highest intensity values (Extended Data Fig. 5b). Although these experiments are not designed to be very quantitative, they suggest that phospholipids are the predominant lipid components present in hemocytes, which is consistent with a previous study of hemocyte lipids^31^. Other lipid classes, such as carnitines (CAR), cholesteryl esters (CE), and diacylglycerols (DAG), display relatively lower intensity signals, suggesting their lesser abundance in comparison to phospholipids (Extended Data Fig. 5b). CE species, which are a major component of LDL^46,47^, displayed a broad intensity distribution in head hemocytes. Likely because of the high sensitivity of MRM-screening approaches, we successfully detected CE in head hemocytes, while prior studies did not^31^. We hypothesize CE results from LDL uptake by hemocytes from glia and is processed when they leave the head cavity (Extended Data Fig. 5b-c). We further focused on the LDL uptake function of Eater in hemocytes.

In previous *in vitro* experiments, the domains 1-199 of the Eater protein were found to interact with acetylated LDL or oxidized LDL^37^. To test the affinity of hemocyte expressed Eater for acetylated or oxidized LDL versus neutral LDL, we employed an *ex vivo* system where wild-type or *eater* mutant hemocytes were cultured with neutral, acetylated, or oxidized LDL. Consistent with prior studies^37^, neither wild-type nor *eater* mutant hemocytes exhibited any affinity for neutral LDL (Extended Data Fig. 5d-e). However, wild-type hemocytes bound more oxidized LDL (Extended Data Fig. 5f-g) and acetylated LDL (Fig. 5a-b) than *eater* mutants. Interestingly, most of the acetylated LDL remained outside the hemocyte (Fig. 5a-b), while oxidized LDL was observed intracellularly (Extended Data Fig. 5f-g). This suggests that hemocytes have different affinities or uptake rates for these modified forms of LDL, with faster uptake of oxidized LDL. Oxidized LDL is predominantly taken up by the CD36 homolog *croquemort* (Crq) in *Drosophila*^30^, but knockdown of *crq* in Hml+ hemocytes or *crq* mutants did not show a strong sleep phenotype (Extended Data Fig. 2a, 5h). On the other hand, hemocyte Eater affects sleep and affects uptake of both oxidized and acetylated lipids.

**Figure 5.**
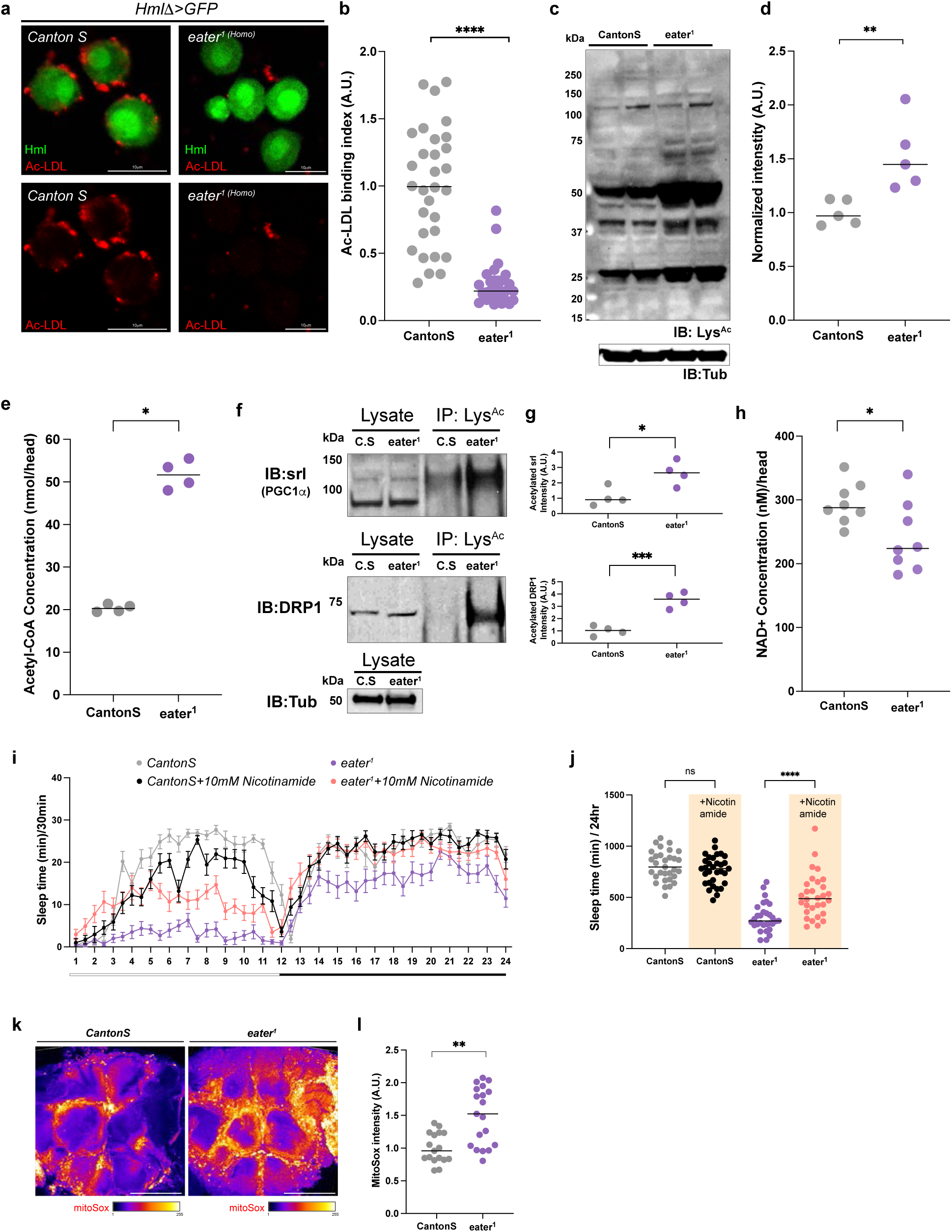
Protein acetylation is increased in the eater mutant brain. **a-b.** *Ex vivo* culture of hemocytes with acetylated LDL. Acetylated LDL (red) was observed on the surface of wild-type hemocytes (left, green, *Hml∆>GFP/+*) but not on *eater* mutant hemocytes (right, green, *Hml∆>GFP/+; eater*^1^) **(a)**. Quantification of the bound acetylated LDL on hemocytes. Bounded amount was normalized to *CantonS* hemocytes **(b)**. Mann-Whitney test was performed for data analysis. **c-d.** Comparison of acetylated proteins in the wild-type (*CantonS*) and *eater* mutant (*eater*^1^) fly brain. The *eater* mutant (*eater*^1^) brain showed an increase in acetylated proteins compared to wild-type (*CantonS*) **(c)**. Quantification of acetylated proteins from the western blot **(d)**. IB: immunoblot. Lys^AC^: acetylated lysine. Tub: alpha tubulin. Mann-Whitney test was performed for data analysis. **e.** Measurement of Acetyl-CoA levels in *eater* mutant (*eater*^1^) and wild-type (*CantonS*) brains. Mann-Whitney test was performed for data analysis. **f-g.** Immunoprecipitation of acetylated proteins in wild-type or *eater* mutant brain. The expression of Srl in the wild-type (*CantonS*, C.S) or *eater* mutant (*eater*^1^) is similar in the lysate but more acetylated-Srl was detected in the *eater* mutant (*eater*^1^) (top). Similarly, DRP1 expression was consistent between wild-type (*CantonS*, C.S) and *eater* mutant (*eater*^1^) lysates, but more acetylated-DRP1 was detected in *eater* mutants (*eater*^1^) (bottom) **(f)**. This was confirmed by quantification of acetylated Srl (top) or DRP1 (bottom) levels in wild-type (*CantonS*, C.S) and *eater* mutant flies (*eater^1^*) (**g**). IB: immunoblot. IP: Immunoprecipitation. Lys^AC^: acetylated lysine. Tub: alpha tubulin. Unpaired t test was performed for data analysis. **h.** Measurement of NAD+ levels in *eater* mutant (*eater^1^*) and wild-type (*CantonS*) brains. Unpaired t test was performed for data analysis. **i-j.** Sleep analysis in female wild-type and *eater* mutant with or without 10mM nicotinamide supplementation. The graph represents sleep in wild-type (*CantonS*) and *eater* mutants (*eater^1^*) with or without 10mM nicotinamide **(i)**. Quantification of the sleep amount from (i) in **(j)**. Orange shade in (j) represents feeding of 10mM nicotinamide. Tukey’s multiple comparisons test was performed for data analysis. **k-l.** Staining of reactive oxygen species (ROS) using MitoSox in the fly brain. Compared to wild-type *(CantonS), eater* mutant *(eater^1^)* has more MitoSox incorporation in the brain **(k)**. Mann-Whitney test was performed for data analysis. Quantification of the MitoSox intensity from (k) in **(l)**. ns: not significant (p>0.01); *p<0.1; **p<0.01, ***p<0.001. ****p<0.0001. Bars in graphs: the median. White scale bar, 100 μm unless otherwise indicated. White and black bar below the graph in (i) represents day (white) and night (black).

We considered the possibility that acetylated lipoproteins, rather than acetylated lipids per se, are targeted by Eater. Lipid droplet transfer is typically mediated by apolipoproteins^48–50^, one such being GLaz, the *Drosophila* ortholog of apolipoprotein E/D (ApoE/D). GLaz is known to be acetylated and, interestingly, knockdown of GLaz decreases sleep in flies^10,51^. However, immunoprecipitation assays indicated that levels of acetylated GLaz are similar between wild-type flies and *eater* mutants (Extended Data Fig. 5i-j), suggesting that it does not contribute to the *eater* phenotype. However, *eater* mutants exhibit an overall increase in acetylated proteins compared to wild-type (Fig. 5c-d).

### Loss of eater results in dysregulated energy metabolism in glia, memory defects and shorter lifespan

Numerous proteins undergo acetylation, and this modification is conserved across a wide range of species, from nematodes to humans^52,53^. Acetylation regulates various cellular processes, including mitochondrial metabolism, protein translation, protein folding, and DNA packaging^54^. And it can be catalyzed by enzymes such as histone acetyltransferases (HATs), or it can occur non-enzymatically when Acetyl-CoA levels are elevated in the cell^52^.

Given the increase of acetylated proteins in *eater* mutants (Fig. 5c-d), we investigated whether the levels of Acetyl-CoA were elevated compared to wild-type flies. Notably, the concentration of Acetyl-CoA in *eater* mutants was more than twice that of wild-type flies (Fig. 5e). We sought to identify candidate proteins that may be targeted by the high Acetyl-CoA, and so considered PGC1α and DRP1, which regulate mitochondrial biogenesis and mitochondrial fission, respectively, and whose acetylation affects proper mitochondrial activity^55,56^. To determine whether PGC1α or DRP1 is more acetylated in the *eater* mutant brain, we immunoprecipitated acetylated lysine and immunoblotted for the PGC1α and DRP1 proteins (Fig. 5f-g). We found that acetylation of both spargel (srl; *Drosophila* homolog of PGC1α) and DRP1 is increased in *eater* mutants (Fig. 5f-g). As compromised mitochondrial activity can affect NAD levels, we measured these in *eater* mutants and found that NAD+ and NADH levels were lower than in the controls (Fig. 5h, Extended Data Fig. 5k).

To determine whether reducing acetylation could rescue sleep in *eater* mutants, we overexpressed the deacetylase enzyme sirtuin^57^ in glia; however, this intervention did not restore total sleep in *eater* mutants (Extended Data Fig. 5l). Since sirtuin activity depends on NAD+, the depleted NAD+ levels in *eater* mutants may explain the lack of rescue by sirtuin overexpression. Indeed, supplementing fly food with 10mM nicotinamide, a precursor for NAD synthesis, which can restore NAD+ levels and thereby enhance endogenous deacetylase activity, partially rescued the *eater* sleep phenotype. Feeding nicotinamide had no effect on sleep in wild-type flies (Fig. 5i-j).

Given the acetylation of key mitochondrial proteins, we assessed mitochondrial integrity by measuring ROS. As a first step, we visualized ROS in the brain using two different fluorescent probes, MitoSox and DHE^58^ (Extended Data Fig. 5m). Because we were particularly interested in cortex glia, we co-localized with a cortex glia marker and compared ROS levels between wild-type and *eater* mutant flies (Extended Data Fig. 5m). *eater* mutants exhibited ROS levels that were 1.5 times higher than those observed in wild-type flies (Fig. 5k-l). However, no cell death was observed in the brain (Extended Data Fig. 5n). Altogether, our results suggest that when lipids are not cleared from glial cells by hemocytes, the resulting lipid accumulation in glia leads to metabolic stress, including increased Acetyl-CoA, reduced NAD+, mitochondrial dysfunction caused by DRP1 or PGC1α acetylation and elevated ROS levels. This metabolic stress likely contributes to sleep loss in *eater* mutant flies.

The reduced sleep and dysregulated metabolic processes in *eater* mutants led us to ask whether memory and lifespan were affected. As seen in some other sleep mutants^59,60^, *eater* mutant flies exhibited deficits in both short-term and long-term memory (Fig. 6a) and shorter lifespan compared to wild-type flies (Fig. 6b).

**Figure 6.**
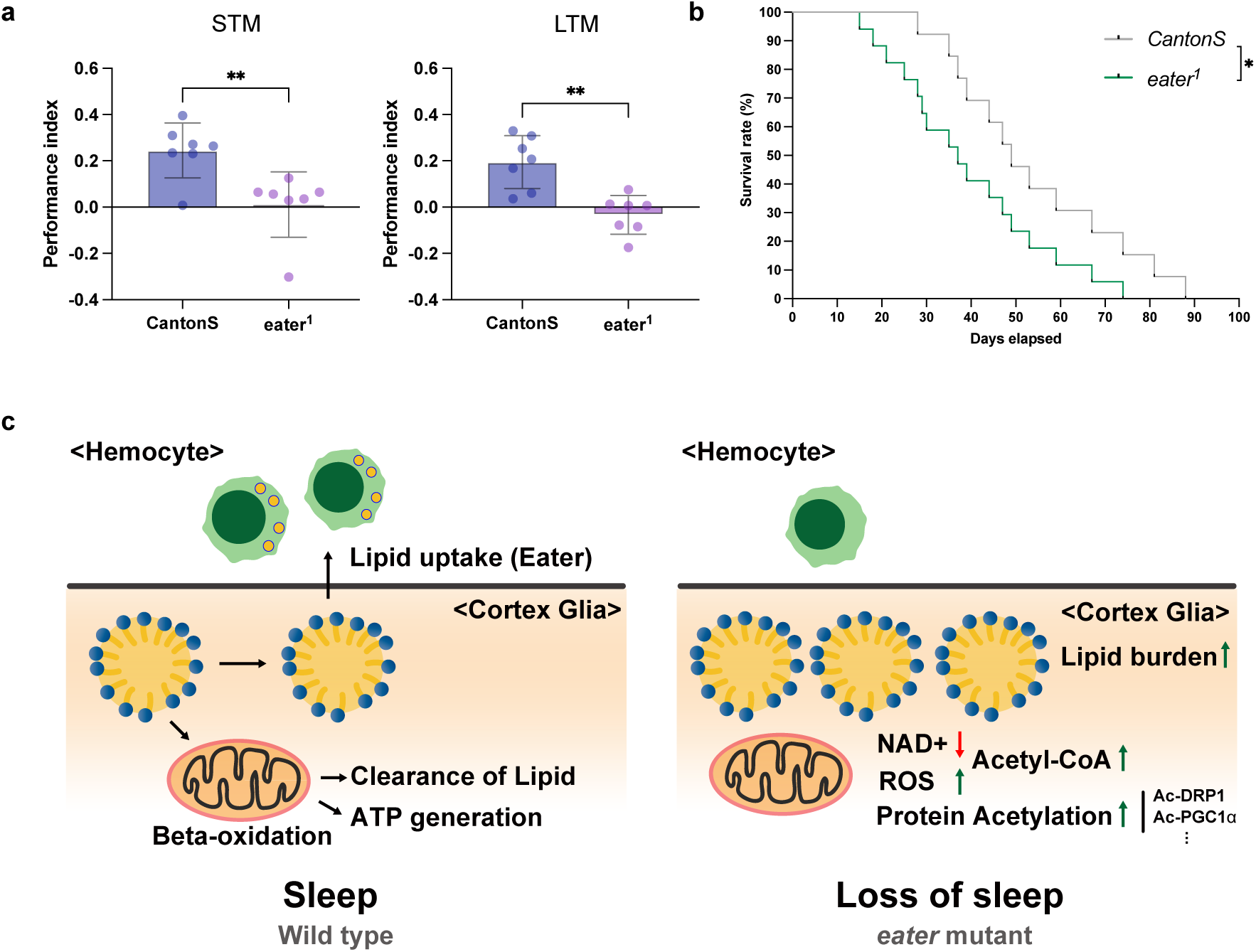
eater mutants have memory deficits and reduced lifespan. a.Measurement of short-term memory (STM, left) or long-term memory (LTM, right) in wild-type (*CantonS*) or *eater* mutants (*eater^1^*). *eater* mutants (*eater^1^*) exhibited impairments in both STM and LTM. Unpaired t test was performed for data analysis. **b.** Comparison of lifespan between wild-type (*CantonS*) and *eater* mutant (*eater^1^*) flies. *eater* mutants showed a reduced lifespan relative to wild-type. Log-rank (Mantel-Cox) test was performed for data analysis. **c.** Schematic illustration of hemocyte-glia interaction during sleep. Lipid droplets can be eliminated in two distinct pathways. One pathway involves lipid catabolism through beta-oxidation in the cortex glia. The other pathway involves the uptake of lipid droplets from the cortex glia by hemocytes via Eater. When hemocyte-mediated lipid uptake is disrupted, lipid droplets accumulate in cortex glia, leading to increased protein acetylation and ROS levels while reducing NAD+ levels. This metabolic regulation between glia and hemocytes is crucial for maintaining proper brain lipid metabolism. Sleep promotes this metabolic regulation and is reduced when the process is disrupted. ns: not significant (p>0.01); *p<0.1; **p<0.01, ***p<0.001. ****p<0.0001. Bars in graphs: the median.

## Discussion

We show here that hemocytes are recruited to the brain during periods of increased sleep and they clear lipids via the Eater protein. If Eater function is impaired, glia accumulate more lipids, and the lipid burden induces metabolic stress with an increase in protein acetylation. This causes mitochondrial dysfunction and metabolic imbalance in the brain (Fig. 6c). In addition to disrupted sleep, memory is impaired and lifespan is shortened. These findings highlight a critical role of brain-periphery interaction, specifically glia-hemocyte lipid transfer, in maintaining brain metabolic health during sleep.

Most studies of immune-sleep interaction have focused on active immune states like inflammation, specific disease, or sleep deprived conditions^61^. In our study, we aimed to investigate the interaction between the immune system and sleep in normal daily conditions, where the immune system is not active. And we focused on the role of immune cells. To achieve this, we used the simple model organism *Drosophila* and found that macrophage-like immune cells in the circulation track to the brain during sleep. We then screened genes expressed in these cells, for their effects on sleep. Through this screening, we identified the gene ‘*eater’*, which encodes a protein with 32 EGF-like repeats that is involved in cell-cell adhesion, LDL uptake, and phagocytosis of gram-positive bacteria^37^. *eater* mutant flies show reduced total sleep and increased sleep fragmentation (Fig. 2a-b) along with memory defects and reduced life span (Fig. 6a-b). We were able to completely rescue sleep as well as adhesion and lipid uptake phenotypes of *eater* mutants by expression of *eater* in hemocytes (Fig. 2c-d).

We find that hemocytes take up lipids from brain cortex glia via the Eater receptor. The lipids taken up are likely those that are transferred to cortex glia from neurons to prevent wake-induced damage to neuronal mitochondria^10^. Thus, the whole process relieves oxidative burden on the brain, which is supported by the increased oxidation seen in the absence of Eater. While transporters that mediate neuron-glia transfer have been identified^10,51^, how exactly lipids are transferred from cortex glia to hemocytes is unclear. Our data confirm physical contact between hemocytes, and glial cells, supporting direct hemocyte-glia interactions. Given our data showing direct contacts between BBB glia and hemocytes, we speculate that LDs from cortex glia are transferred through the BBB to hemocytes; direct contacts between hemocytes and cortex glia may also occur, but would require validation. While LD accumulation predominantly occurs in cortex glia, we note that other glial subpopulations, in particular astrocytes, also transfer LSD2::GFP-labeled lipids to hemocytes. It is possible that astrocytes process lipids without accumulating LDs or, alternatively, that they transfer droplets or fatty acids via lipoprotein particles to other glia^45^. Ultimately, many of these lipids end up in hemocytes. However, some are likely also processed in glia via beta oxidation, thereby generating energy. Likewise, we speculate that the lipids transported in hemocytes are processed, either within the hemocytes themselves or in the fat body. Interestingly, specific lipid binding/processing molecules have been implicated in the regulation of sleep^62,63^.

Eater is necessary for the transfer of lipids associated with LDs that predominantly accumulate in cortex glia; without *eater* more LDs accumulate in the brain, and fewer in hemocytes, highlighting the importance of *eater* for clearing glia lipids. In addition to increased LDs, *eater* mutant brains had higher levels of acetylated proteins, increased Acetyl-CoA levels, and reduced NAD+ levels. Although the sequence of these changes remains uncertain, we propose that loss of lipid uptake by Eater leads to an accumulation of LDs, triggering metabolic stress characterized by elevated Acetyl-CoA, increased acetylation of key mitochondrial proteins, and impaired mitochondrial function. Increased Acetyl-CoA levels also indicate less beta-oxidation and lower energy production, which further compromises the ability of mitochondria to oxidize fatty acids. This fuels a vicious cycle of metabolic stress, oxidative damage and LD accumulation. The consequently reduced levels of NAD+ may further contribute to increased acetylation by impairing NAD-dependent deacetylase enzymes, such as sirtuins^57^. However, we cannot exclude the possibility that increased Acetyl-CoA levels are a consequence of reduced uptake of acetylated lipids by Eater. Or that lower NAD is an early outcome of the metabolic stress and that it accounts for higher acetylation of DRP1 and PGC1α. Acetylation of PGC1α is known to inhibit its own functions^64^ while acetylation of DRP1 increases its activity but induces metabolic stress and cellular dysfunction^56^. Haynes et al. previously demonstrated that knock-down of *Drp1* in neurons or glia decreases sleep, as does knockdown of beta-oxidation-related genes, such as *Mcad*^10^. Similar impairments of mitochondrial function likely result from acetylation of DRP1 and PGC1α, the latter being a transcription factor that promotes mitochondrial biogenesis and the expression of beta-oxidation-related genes^65^.

Protein acetylation (beyond histones) has been mostly studied in the context of metabolic syndromes like alcoholic liver disease, high fat diet, or atherosclerosis^66^, with less known about its role in other biological processes. We found that *eater* mutants have elevated acetylated proteins and lower NAD+ levels. Recent research on short sleep mutants has identified decreased NAD+ levels in the brain^67^ but, to our knowledge, protein acetylation has not been examined in the context of sleep regulation. These findings suggest that the interplay between NAD+ levels and protein acetylation in the brain may play a critical role in sleep control and function.

Currently we do not know what regulates hemocyte numbers in the head cavity during the wake-sleep cycle. Previous work with pupae found that heart rhythms remain constant during the day and night^68^, suggesting that changes in hemocyte numbers are unlikely to be due to heartbeat fluctuations but may instead result from changes in hemolymph flow driven by fly movement during wake versus sleep. However, hemocyte numbers change within the sleep state, suggesting that they are influenced by other factors. One possibility is ROS production by glial cells. ROS levels in the glial mitochondria increase during wakefulness and decrease with sleep^10^. In mammals, ROS promotes leukocyte adhesion and penetration, and it also serves as an adhesion cue for hemocytes in flies^69,70^. Therefore, fluctuations in hemocyte numbers could be linked to ROS levels, although this would predict higher adhesion to the brain at the end of wake rather than the middle of the night.

While brain-periphery interactions are currently receiving attention, the role we report here for hemocytes is unprecedented. Our findings suggest that oxidated and acetylated lipids need to be removed from the brain by hemocytes to prevent oxidative damage and preserve the integrity of brain mitochondria. In mammals, microglia are key glial cell types that take up lipids from neurons, and are particularly important in the context of neurodegeneration^71–73^. Since *Drosophila* lack microglia, circulating hemocytes may serve an analogous function, acting as intermediaries for lipid uptake and transport/storage and combating stress by accumulating LDs. Importantly, we find that this is a sleep-dependent process, indicating that peripheral blood cells perform a function of sleep. This represents a major shift in paradigm, given the previous brain-centric view of sleep and even the current brain-focused hypotheses for why we sleep^2,74^.

## Material and Method

### Drosophila strains and fly husbandry

All flies for experiments were maintained at 25°C in a 12h-12h light-dark cycle, except for the temperature sensitive Gal80 experiment in which Gal80 flies were kept at 18°C until the experiment and activated Gal4 at 30°C for two days. For the gene switch experiment, 500 μM mifepristone (RU486, sigma, M8046) was added to sucrose/agar food. For the RU486 control food, same amount of 80% ethanol was added to sucrose/agar food. The following *Drosophila* stocks were used in this study: CantonS (Sehgal Lab stock), HmlΔ-LexA LexAop-mCherry (J.Shim), HmlΔ-Gal4 UAS-EGFP (BL30139, BL30140), Srp-Hemo-mCherry (BL78358, BL78359, BL78362, BL78363), 9-137-Gal4 (Sehgal Lab stock), NP2222-Gal4 (Sehgal Lab stock), R85G01-LexA (BL54285), R54C07-LexA (BL61562), Srp-Gal4 (L.Waltzer), Srp-Hemo-Gal4^DBD^ (I. Evans), Srp-Hemo-Gal4^AD^ (Iwan Evans), eater-dsRed (U.Banerjee), Ppn-Gal4 (BL77733), NP6293-Gal4 (Sehgal Lab stock), moody-Gal4 (Sehgal Lab stock), Repo-GeneSwitch (Sehgal Lab stock), UAS-CD4::GRASP (BL58755), eater^1^ (BL68388), UAS-eater (BL36325), eater RNAi (BL25863, V4301), UAS-mCD8GFP (BL5137), UAS-mCD8::RFP, LexAop-mCD8::GFP (BL58754) UAS-LD::GFP (M. Welte), UAS-GFP::LSD2 (M. Welte), UAS-Sirt1 (BL44216), GLaz-GFSTF (BL60526), UAS-h.MEGF11(BL78460), MZ0709-Gal4 (M.Freeman), Alrm-Gal4 (M.Freeman), Repo-Gal4 (L. Griffith), Repo-LexA (M.Freeman), put RNAi (BL35195), Lsd-1 RNAi (V30884), NimC3 RNAi (V22920), GLaz RNAi (V4806), Lsd-2 RNAi (V40734), UAS-PvrDN (BL58431), Nplp2 RNAi (BL54041), NimC1 RNAi (BL25787), NimC4 RNAi (BL61866), NimB1 RNAi (BL55937), NimC2 RNAi (BL25960), Smox RNAi (BL41670), PGRP-LC RNAi (BL33383), drpr RNAi (BL67034), NimB4 RNAi (BL55963), Apoltp RNAi (BL 51937), Nim A RNAi (V104204), Col4a1 RNAi (BL44520), NimB2 RNAi (BL62289), NimB5 RNAi (V15758), babo RNAi (BL25933), Karl RNAi (V9446), LpR2 RNAi (BL31150), apolpp RNAi (BL28946), Acsl RNAi (V3222), LpR1 RNAi (BL27249), crq RNAi (BL40831), NimB3 RNAi (V330502).

### Sleep recording

For the fly sleep recording, mated 5-7 days old flies were loaded into glass tubes containing 5% sucrose with 2% agarose. At least two days after loading into the monitors, sleep was analyzed for three days. Single beam monitors were used for RNAi screening and the sleep deprivation experiment, but other sleep recordings were performed with multibeam monitors. Sleep was defined as failure of the fly to cross the red beam in the monitor for 5 or more minutes, analysis of data was performed with an in-lab built code^75^ like previous.

### Sleep deprivation

We used mechanical sleep deprivation. To achieve deprivation, flies in single beam monitors were fixed to a vortex machine and randomly shaken for 2 seconds every 20 seconds over a 12 hour period (From ZT12-24). 6 hours of sleep after sleep deprivation was compared to sleep on the pre-deprivation day at the same ZT time (ZT 0-6) to estimate rebound sleep. Analysis of percent sleep gain was as described previously^76^. In short, 6 hours of daytime sleep on the day prior to deprivation was subtracted from the 6 hours after deprivation (Sleep gain). Similarly, sleep loss was calculated by subtracting sleep during deprivation to nighttime sleep a day before the deprivation (Sleep loss). Percentage of sleep gain was calculated by amount of sleep gain relative to amount of sleep loss.

### Fly tissue clearing

For visualizing circulating hemocytes, we optimized two different protocols^77,78^. Heads of *HmlΔ-LexA LexAop-mCheery* flies were cut with a micro scissor and fixed with 4% paraformaldehyde for 4 hours at room temperature with a rotation. After the fixation, heads were incubated with 100% methanol at the 4°C for overnight. Methanol was removed the following day and heads were incubated with BABB solution (2:1 ratio of benzyl benzoate and benzyl alcohol) at least 6 hours. After the removing the BABB solution, heads were mounted on the slide glass with VECTASHIELD solution without DAPI. Imaging was performed right after the mounting and 408 excitation laser was used for auto-fluorescent signal.

### Immunohistochemistry

Fly brains were fixed in a 4% paraformaldehyde solution and washed three times using 0.4% PBS TritonX-100. After the three washes, samples were blocked using 10% NGS for 30 min at 25°C. Samples were incubated with the desired primary antibodies overnight at 4°C and then washed three times using 0.4% PBS TritonX-100 (PBST). Samples were incubated with secondary antibodies (Life Tech, A32723, A32740, A32742, A32731, A21236) diluted 1:250 for 2 h. Samples were then washed three times with 0.4% PBST. After the washing, samples were rinsed and kept in VECTASHIELD until they were mounted on glass slides. For hemocytes count, we did not remove air sacs to maximize the number of hemocytes. The following primary antibodies were used: ɑ-NimC1 (Gift from I. Ando, 1:100), ɑ-brp (DHSB, nc82, 1:100), ɑ-Repo (DSHB 8D12, 1:100), ɑ-cleaved dcp1(Cell signaling, 9578S, 1:100), Oil-Red O (Sigma, O9755), BODIPY 493/503 (Fisher, D3922, 1:1000). Images were obtained with the Leica Stellaris STED confocal microscope. For the hemocyte counts, we used 3D object counter in Image J software.

### Staining of lipid droplets using Oil-Red O

Fly brains were dissected and fixed as for immunohistochemistry. After the fixation, brains were washed three times with 0.4% PBS TritonX-100 and kept in 0.4% PBST overnight at 4°C. If a sample needed primary antibody incubation, it was blocked with 10% NGS in 0.4% PBS TritonX-100 for 30 min and then kept in diluted antibody with 0.4% PBST overnight at 4°C. The following day, Oil-Red O solution (0.1g/20ml of isopropanol; Sigma, O0625) was prepared in 0.4% PBST as a two to three ratio. If the sample was not stained with primary antibody, sample was incubated with Oil-Red O solution 10 minutes and washed with distilled water five times for 5 minutes each. If samples were stained with primary antibody, samples were washed and treated with secondary antibody and after the secondary antibody incubation Oil-Red O solution was treated. Lastly, samples were rinsed with PBS and mounted in VECTASHIELD until they were mounted on glass slides. Images were obtained with the Leica Stellaris STED confocal microscope.

### Quantification of Lipid droplets in the brain

To count lipid droplets, images were analyzed with a custom ImageJ macro. For each slice in the stack, the BioVoxxel toolbox was used to subtract background noise using the convoluted background subtraction method with a mean convolution filter of 3 radius. Once the background was subtracted, the image was duplicated, and one of the copies was converted into a mask that contained only the brain region. Inside the masked area, lipid droplets were counted using the Analyze Particles tool, defining a particle size of 2-250 and circularity of 0.4-1. Reported results are the sum of particles in the whole stack. The macro is publicly available at https://github.com/CamiloGuevaraEsp/lipid_droplets.

### Immunoprecipitation and western blot

In a 15ml tube, at least 100 flies were collected for immunoprecipitation, or 20 flies were used for western blot. Flies were frozen on dry ice for 10 min. After freezing, flies were vortexed 10-20 seconds three times to shake off heads. Flies were then poured into a sieve that only allows passage of heads. Fly heads were homogenized at 25Hz for 2 minutes in TissueLyser II (Qiagen) in 100ul of lysis buffer for immunoprecipitation (250mM Tris-HCl (pH 7.5), 250mM NaCl, 1.5M Sucrose, 1% TritonX-100, protease inhibitor cocktail) or for western blot (RIPA buffer, Lifetech, 89901) with a 5mm stainless steel bead (Qiagen, 69989) in round bottom tubes (USA Scientific, 1620-2700). Homogenized samples were transferred to 1.7ml microcentrifuge tubes and spun for 14000 rpm, 10 minutes at 4°C. Supernatants were used for the experiment. For immunoprecipitation, protein A/G magnetic agarose beads (Fisher, 78609) were used for antibody conjugation. Antibody conjugation to the bead or incubation of antibody with samples was performed in the cold room overnight. Samples were run in a 4-12% premade gel (Life Tech, NP0322). Following antibodies were used for experiment. ɑ-Acetylated lysine-Mouse (Life Tech, MA12021, 1:1000), ɑ-Acetylated lysine-rabbit (Cell signaling, 9441S, 1:1000), ɑ-DRP1 (Gift from L.Fisher, 1:1000), ɑ-SRL (Gift from A.Duttaroy, 1:1000), ɑ-alpha Tubulin (DSHB, 12G10, 1:1000), ɑ-FLAG (Sigma, F3165, 1:2000), ɑ-mouse-HRP (Jackson Immuno, 715-035-151, 1:2000), ɑ-rabbit-HRP (Jackson Immuno, 715-035-152, 1:2000) and HRP signal was obtained with ECL substrate (Life Tech, 32209).

### Flow cytometry

At least 100 *Hml∆-Gal4 UAS-EGFP* fly heads were cut with a micro scissor under the microscope and kept in the round bottom tube with 1000ul of ice-cold Schneider’s medium (Life Tech, 21720024) until the cutting was finished. With a metal bead, fly heads were homogenized with TissueLyser II (Qiagen) at 25Hz for 2 min. Samples were centrifuged at 6000 rpm, for 5 min at 4°C. Supernatant was discarded and pellets were treated with 37°C pre-warmed 100ul of Collagenase Type C (100mg/ml, Worthington-Biochem, LS004140), 380ul of PBS, 20ul of Dispase II (100mg/ml, Sigma, D4693). Samples were incubated on a rotator for 15 min and pipetted with a 200ul pipet every 5 min. 200ul of ice-cold PBS was added to the sample, which was transferred to a 1.7ml microcentrifuge tube. Samples were spun at 6000 rpm, for 5 min at 4°C. Supernatant was discarded and pellets were resuspended in 500ul of cold Schneider’s medium. 0.5 ul of DAPI (1mg/ml) was added and after short vortexing, samples were spun at 6000 rpm for 5 min at 4°C. Again, supernatant was discarded, and samples were resuspended in 600ul of Schneider’s medium. Using a 40μm strainer (Sigma, BAH136800040), debris or clumps were removed, and samples were transferred to a 5ml FACS sorting tube. We used Aria FACS sorter (BD Biosciences) with 100 μm nozzle. Usually, 100 fly heads give approximately 400 GFP+ hemocytes after sorting.

### Hemocytes transfer

At least 30 larvae were dissected in Schneider’s medium (Life Tech, 21720024) and kept on ice during the dissection. Samples were spun at 6000 rpm, for 5 min at 4°C. Hemocytes were resuspended with 100ul of PBS for a cell density of 100-150 cells per microliter. Using the microneedle, 2-3 ul of hemocytes were injected into the fly thorax. *HmlΔ-LexA LexAop-mCherry* hemocytes were used for validating hemocyte transfer.

### In vitro hemocyte culture

Larvae were dissected in 15ul of Schneider’s medium (Life Tech, 21720024). Hemocytes were transferred to Schneider’s medium containing Dil-labeled neutral, oxidized, or acetylated LDL (1:100 dilution, Life Tech, L3482, L34358, L3484) in the tube. Hemocytes were incubated on teflon printed microscopic slides (immune-cell, 61-100-17) for 2 hours in the cold room. After the 2 hours, hemocytes were fixed with 4% paraformaldehyde and washed three times with 0.4%PBST. Hemocytes were kept in VECTASHIELD with DAPI before the imaging and images were obtained with Leica Stellaris STED confocal microscope.

### NAM food feeding

Flies were raised on normal food for 5 days after eclosion and then transferred to sleep recording glass tubes which contain 5% sucrose and 2% agarose food with or without 10mM nicotinamide (Sigma, 72345).

### Gaboxadol feeding

Gaboxadol hydrochloride (Sigma, T101) was dissolved in the normal fly food at a 2mM concentration. Flies were kept in this for two days.

### Acetyl CoA measurement

To measure Acetyl-CoA in the fly head, 10 female flies were collected in the 15ml falcon tube and frozen on dry ice for 10 minutes. After freezing, flies were vortexed 10-20 seconds three times to shake off heads. Flies were then poured into a sieve that only allows passage of heads. 10 fly heads were homogenized with 25Hz for 2 minutes in TissueLyser II (Qiagen) in the extraction buffer from the Acetyl-CoA colorimetric assay kit (Elabscience, E-BC-K652-M). After the homogenization, all the procedures followed manufacture’s instructions from the kit.

### NAD^+^/NADH measurement

To measure NAD^+^/NADH in the fly head, 10 male and female *CantonS* and *eater*^1^ mutant flies were collected around ZT6 in 1.5ml tubes then flash frozen on dry ice. After freezing, flies were vortexed, and 10 heads were collected then placed in a 2ml tube with a metal beads and 1 mL of lysis buffer (1:1 PBS: Extraction buffer (10% DTAB 0.2 NaOH)). The heads were homogenized using the TissueLyser II (Qiagen) at 25 Hz for 2 min. The homogenized liquid was passed through homogenizer tubes (Invitrogen, 12183-026) in a centrifuge for 5 min at 15,000 rpm at 4°C. In order to measure NAD^+^ and NADH individually, 200 μL of lysate was added to separate 1.5ml tubes; 100 mL of 4M HCl was added to the NAD^+^ tube, and both were heated for 15min at 60°C. After incubation at room temperature for 10 minutes, 100 μL of 0.5 M Trizma base buffer was added to the NAD^+^ tube, and 200 μL neutralization buffer (a 1:1 mixture of 0.5 M Trizma: 0.4 M HCl) was added to the NADH tube. NAD^+^ samples were diluted 1:5 and NADH samples were diluted 1:1 using dilution buffer. Samples and standards were prepared 1:1 with NAD^+^/NADH-Glo detection reagent (Promega, G9071) according to manufacturer’s instructions, seeded onto a 384-well plate, and measured using a BioTek Cytation 5 imaging reader and the accompanying Gen5 v3.12 software. Individual data points are the mean of 3 technical replicates.

### MitoSox Staining

To stain the fly brain with MitoSox (Fisher, M36008), 10 fly brains were dissected in Schneider’s medium and kept in medium until the dissection was finished. Brains were transferred to Schneider’s medium with MitoSox dye (Final concentration 5μM) and incubated for 10 minutes with rotation at room temperature. After the incubation, brains were washed with Schneider’s medium three times for three minutes with rotation. Brains were fixed with 4% paraformaldehyde for two minutes and rinsed with PBS. MitoSox signal was imaged right after the mounting with the VECTA SHIELD using the Leica Stellaris STED confocal microscope.

### Memory experiment

Memory experiment was as published^59^. In brief, 100 flies, 3-6 day old and of mixed sex, from the same genotype were starved in agarose for 18 hours. Following day, flies were trained at 25°C in 70% humidity in a small chamber containing 1.5M sucrose or water soaked whatman paper with odors (1:200 ratio of MCH or 1:80 ratio of OCT in the paraffin oil) for 2 minutes under red light. After the training, flies were located in the bidirectional choice apparatus which has an odor in each end. To remove the bias coming from the odor, appetitive training was reciprocally performed. For long-term memory, trained flies were kept again in the agarose food for 18 hours and memory experiment was performed without re-training.

### Lifespan experiment

Age matched 300 female or male flies were collected for 5 days and transferred to vials of 30 flies each. Every two-or three-days, flies were flipped into new vials and the number of flies was counted.

### Hemocyte sample Preparation for MS Analysis

From the *Hml∆-Gal4 UAS-EGFP* fly, hemocytes were sorted from the flow cytometry and 4,000 GFP+ cells were sorted to 1.7ml microcentrifuge tube. GFP+ cells were spin downed with 6000 rpm, for 5 min at 4°C and pellets were kept in the -80°C until the lipid extraction. Lipid extraction from fly hemocyte was performed using a modified Bligh & Dyer method^79^. Briefly, frozen cell pellets were thawed at room temperature for 10 minutes before the addition of 200 µL of ultrapure water to facilitate cell lysis. This was followed by the addition of 450 µL of methanol and 250 µL of HPLC-grade chloroform. During extraction, 5 µL of 1 µg/mL of an internal standard mixture comprised of equiSPLASH (Avanti Polar Lipids, #330731), FA 16:0-d2 (Cayman Chemical), and CAR 14:0-d3 (Cayman Chemical) was added to each sample. The samples were vortexed for 10 seconds to form a single-phase solution and incubated at 4°C for 15 minutes. Subsequently, 250 µL of ultrapure water and 250 µL of chloroform were added, inducing phase separation. The samples were then centrifuged at 16,000g for 10 minutes. The organic phase, containing the extracted lipids, was carefully transferred to fresh tubes and evaporated using a vacuum concentrator to obtain dried lipid extracts.

The dried lipid extracts were reconstituted in 200 µL of a 3:1 methanol:chloroform (MeOH:CHCl₃) containing 10 mM ammonium formate. Following reconstitution, all samples were analyzed using multiple reaction monitoring (MRM) methods. An injection solvent containing 0.02 µg/mL EquiSPLASH® (Avanti Polar Lipids, #330731) was used as a quality control sample to monitor peak stability over time.

### Unbiased Lipidomics via Multiple Reaction Monitoring (MRM)-Profiling

Lipidomic analyses were performed using an Agilent 6495C triple quadrupole mass spectrometer (Santa Clara, CA) coupled to an Agilent 1290 Infinity II LC system with a G7167B autosampler. Samples were introduced into the Agilent Jet Stream (AJS) ion source via direct flow injection (i.e., no chromatographic separation). MS data was acquired for 3 min per injection. For each multiple reaction monitoring (MRM) scan method, 8 μL of the sample was injected. MRM methods were organized into 25 methods on the basis of the ten main lipid classes based on the LipidMaps database, spanning over a total of 3000 individual lipid species. TAGs and DAGs were divided into separate methods based on fatty acid neutral loss residues. Statistical analysis was performed using the EdgeR package^80^. EdgeR uses a generalized linear model to identify differentially expressed lipids. This method was described in detail in previously^45,73,81^. Significant lipids were chosen based on a false discovery rate value <0.1.

## Acknowledgments

The authors thank the members of the Sehgal lab and Chopra lab for helpful discussions. The authors acknowledge the Bloomington, VDRC, DGRC and the DSHB hybridoma bank. The authors thank the following individuals for stocks and reagents: Drs. J.Shim, I. Ando, I.Evans, U.Banerjee, M.Welte, M.Freeman, L.Fisher, A.Duttory. This study was supported by grants from the National Research Foundation (NRF) of Korea (RS-2024-00408937) to B.C. and National Institute of Neurological Disorders and Stroke (NS48471) to A.S. This work is funded, in part, by the National Institutes of Health (NIH) National Center for Advancing Translational Sciences award U18TR004146, ASPIRE Challenge and Reduction-to-Practice awards and AnalytiXIN Fellowship award to G.C. C.E.R. acknowledges the Arnold O. Beckman Postdoctoral Fellowship in Chemical Instrumentation Award Program. We thank Agilent Technologies for their gift of the Triple Quadrupole LC/MS to the Chopra Laboratory. The Purdue University Center for Cancer Research funded by NIH grant P30 CA023168 is also acknowledged. The content is solely the responsibility of the authors and does not necessarily represent the official views of NIH. G.C. is the James Tarpo Jr. and Margaret Tarpo Associate Professor of Chemistry.

## Author contributions

Conceptualization: A.S. and B.C.; Methodology: B.C., D.E.Y., S.K., C.G., C.E.R.,

C.H.B., G.C., and A.S.; Investigation and analysis: B.C., D.E.Y., C.E.R., C.H.B., and S.K.;

Data curation: B.C., D.E.Y., C.E.R., C.H.B., and S.K.; Writing: B.C., D.E.Y., C.G., C.E.R.,

P.S., G.C., and A.S.; Funding acquisition and supervision: G.C. and A.S.

## Competing interests

The authors declare the following competing financial interest(s): G.C. is the Director of the Merck-Purdue Center funded by Merck Sharp & Dohme, a subsidiary of Merck and the co-founder of Meditati Inc. and BrainGnosis Inc. The remaining authors declare no competing interests.

## Data and Materials Availability

All data generated during and/or analyzed in this study are included in this published article and its supplementary information and materials that were newly generated for this study, such as plasmids and fly lines, are available from the Lead Contact upon request.

## Extended Data Figure Legends

**Extended Data Figure 1. Blood cells (hemocytes) circulate in the fly head cavity.**

**a.** Localization of *HmlΔ-LexA*+ hemocytes (*Hml Δ-LexA LexAop-mCherry*, red) in the fly.

Structure of the fly was visualized with a UV laser (Autofluorescence, white).

**b-f.** Localization of the hemocytes near the brain. Visualizing of hemocyte with *Srp-Gal4* (Top left, green), *Srp-hemo-Gal4^DBD^/^AD^* (Top right, green), *Ppn-Gal4* (Bottom left, green), or *eated-dsRed* (Bottom right, red). Brain was visualized with brp (magenta), DAPI (Blue). BBB was visualized with *9-137-Gal4* (bottom right, green) **(b)**. Hemocytes (Hml; green, NimC1; red) are not detected in the ventral nerve cord **(c).** NimC1+ hemocytes (red) are located next to the Perineurial glia (PG, green, *NP6293-Gal4 UAS-mCD8GFP*) **(d)** or Sub-perineurial glia (SPG, green, *moody-Gal4 UAS-mCD8GFP*) **(e)**. 3D reconstruction of Figure 1E shows that NimC1+ (magenta) hemocytes physically interact with perineurial glial cells (green) **(f)**. Cortex glial cells were visualized with RFP (red) (*R85G01-LexA LexAop-mCD8GFP NP2222-Gal4 UAS-mCD8RFP*).

**g-h.** Strong GRASP signals (green) were detected between Repo+ glial cells (magenta) and hemocytes, using *Repo-LexA* and *HmlΔ-Gal4* to drive the two halves of GFP (top). In contrast, little green signal was observed in the *Repo-LexA* GRASP control samples (middle) or *HmlΔ- Gal4* GRASP control samples (bottom). Neuropil was visualized using brp (white) **(g)**. With the use of *SPG-LexA* (*R54C07-LexA)* and *HmlΔ-Gal4,* strong GRASP signals (green; left) were detected but not in *SPG-LexA* GRASP control samples (right). Cell membrane was visualized using phalloidin (magenta) **(h)**. To detect *SPG-LexA* and *HmlΔ-Gal4* GRASP signals, GRASP antibody was used.

DAPI: blue, White scale bar, 100 μm unless it’s not indicated.

**Extended Data Figure 2. eater is required in hemocytes to regulate sleep.**

**a.** Screen for genes regulating sleep in hemocytes using RNAi knockdown with *Hml∆-Gal4*.

The sleep amount in wild-type flies is indicated by a black column with a dotted line. Knockdown of *eater* using two different RNAi lines resulted in decreased daytime or nighttime sleep in male and female (columns with red lines).

**a. b.** Graphs represent sleep in wild-type (*CantonS*) and *eater* knockdown flies (*HmlΔ-Gal4 UAS-GFP*, *eater RNAi*) for both males (top, dark green) and females (bottom, purple). Daytime and nighttime sleep are reduced in males (top, dark green) and mostly night-time in females (bottom, purple). Tukey’s multiple comparisons test was performed for data analysis.

**b.** Quantification of sleep time in *eater* knockdown using a different hemocyte driver (*Srp- Gal4*). Sleep is shown in males (top) and females (bottom). Tukey’s multiple comparisons test was performed for data analysis.

**c.** Quantification of sleep in homozygous or heterozygous *eater* mutant flies (top, dark green, male) (bottom, purple, female). Tukey’s multiple comparisons test was performed for data analysis.

**d.** Hemocyte specific overexpression of *h.MEGF11* in *eater* (*eater*^1^) mutant background.

Graphs represents total sleep time. Dark green dots represent *eater* mutant males. Yellow dots represent hemocyte specific *h.MEGF11* overexpression in the *eater* mutant males (top). Purple dots represent *eater* mutant female. Pink dots represent hemocyte specific *h.MEGF11* overexpression in the *eater* mutant female (bottom). Tukey’s multiple comparisons test was performed for data analysis.

**f.** Comparison of sleep time in *eater* knock-down flies only in the adult stage using the temperature sensitive Gal80 (*HmlΔ-Gal4 UAS-GFP*, *Tub-Gal80^ts^*, *eater RNAi*). Graph on the top left side represents total sleep time before Gal4 activation. Graph in the top middle represents total sleep time during Gal4 activation. Graph on the top right side represents total sleep time following return to de-activating temperature. Red shade indicates Gal4 activating time. Quantification on the bottom. Bars in graphs: the median with SEM. White and black bar below the graph represents day (white) and night (black). Tukey’s multiple comparisons test was performed for data analysis.

**g.** Quantification of circadian periods in wild-type (*CantonS*) or *eater* mutant (*eater*^1^) in dark-dark condition. Mann-Whitney test was performed for data analysis.

ns: not significant (p>0.01); *p<0.1; **p<0.01, ***p<0.001. ****p<0.0001. Bars in graphs: the median.

**Extended Data Figure 3. Hemocyte transfer rescues the eater mutant sleep phenotype.**

**a.** Schematic representation of hemocyte transfer experiment.

**b-c.** Confirmation of larval hemocyte transfer to the adult fly in wild-type **(b)** and *eater* mutants

**(c)**. *Hml-LexA*+ hemocytes transferred from the larvae (*Hml Δ-LexA LexAop-mCherry*, red) were visualized in the fly. Structure of the fly was visualized with UV laser (Autoflourescence, white). Magnified images from the dotted box are on the right side.

Extended Data Figure 4. Hemocytes take up lipids from cortex glia cells through eater.

**a-b.** Detection of Hml+ (green) hemocytes within wild-type (*HmlΔ>GFP/+*) and *eater* knockdown (*HmlΔ>GFP, eater RNAi*) flies **(a)**. Quantification of Hml+ hemocytes near the brain showed reduction in *eater* mutants **(b)**. Mann-Whitney test was performed for data analysis.

**c.** Detection of Hml+ (green) hemocytes within wild-type (*HmlΔ>GFP/+*; top left), *eater* mutant (*HmlΔ>GFP/+, eater*^1^; top right) and *eater* rescue background (*HmlΔ>GFP, UAS- eater/+, eater*^1^; bottom left). Quantification of hemocytes is on the bottom right. Tukey’s multiple comparisons test was performed for data analysis.

**d-e.** Detection of lipid droplets in hemocytes. Oil-Red O lipid staining in hemocytes (red) localized to lipid droplets labeled with GFP (green, *HmlΔ-Gal4/UAS-LD::GFP*) **(d)**. BODIPY (green) positive lipid droplet in the Srp-hemo+ (red) hemocyte **(e)**.

**f-g.** Detection of lipid droplets in the brain with Oil-Red O (gray) in wild-type (*HmlΔ>GFP/+*) and *eater* knockdown (*HmlΔ>GFP, eater RNAi*) **(f)**. Quantification of lipid droplets is shown in **(g)**. Mann-Whitney test was performed for data analysis.

**h-i.** Detection of Oil-Red O positive lipid droplets (red) in the brain with a blood-brain barrier marker in wild-type (top, *9-137-Gal4 UAS-mCD8GFP/+*) or *eater* mutant (bottom, *9-137-Gal4 UAS-mCD8GFP/+, eater*^1^). Pearson’s coefficient of Oil-Red O co-localization with BBB glial cell marker **(i)**. Unpaired t test was performed for data analysis in (i).

**j-k.** Detection of lipid droplets derived from different glial cells in hemocytes. NimC1+ (red) hemocytes were co-localized with LSD2::GFP positive lipid droplets originating from all glia (*Repo-Gal4 UAS-LSD2::GFP)*, astrocyte-like glia (*Alrm-Gal4 UAS-LSD2::GFP*), ensheathing glia (*MZ0709-Gal4 UAS-LSD2::GFP*), and BBB glia (*9-137-Gal4 UAS-LSD2::GFP*) **(j)**.

Quantification of LSD2::GFP+ hemocytes is in **(k)**.

ns: not significant (p>0.01); *p<0.1; **p<0.01, ***p<0.001. ****p<0.0001. Bars in graphs: the median with SD (K). DAPI: blue, White scale bar, 100 μm unless otherwise indicated.

**Extended Data Figure 5. Protein acetylation is increased in the eater mutant brain.**

**a-c.** Analysis of lipid classes in hemocytes. A lipidomic analysis pipeline utilizing Multiple Reaction Monitoring (MRM) profiling with an Agilent 6495C Triple Quadrupole Mass Spectrometer (MS) followed by automated data analysis **(a)**. The data are depicted as a boxplot representation of the intensity distribution of various lipid classes detected in wild-type hemocyte samples (*HmlΔ>GFP/+*, N=3) using multiple reaction monitoring (MRM)-based lipidomics. The full intensity distribution across lipid classes, highlighting variations in signal intensities, is shown (left). On the right is a zoomed-in view providing a more detailed comparison of lipid classes with lower intensity signals (right). Each lipid class is color-coded, and individual data points represent detected lipid species **(b)**. Diacylglycerols (DAGs), triacylglycerols (TAGs), and lipid classes in hemocytes are depicted by MRM intensity distribution. Pie charts represent the relative intensity distribution of DAG species categorized by fatty acid (FA) chain composition across three biological replicates (N1, N2, N3) (top). The relative intensity distribution of TAG species is indicated by FA chain composition, showcasing the diversity of FA chains (middle). Predominant lipid classes detected are indicated by lipid class distribution within the lipidomic profile (bottom). The color legends correspond to the FA chain composition and lipid classes in each panel **(c)**.

**d-g.** *Ex vivo* culture of hemocytes with neutral or oxidized LDL. Neutral LDL (red) were barely observed within or attached to wild-type (left, green, *HmlΔ>GFP/+*) or *eater* mutant (right, green, *HmlΔ>GFP/+, eater*^1^) hemocytes **(d)**. Quantification of the bound neutral LDL on the hemocytes. Bounded amount was normalized to *Canton S* hemocytes **(e)**. Oxidized LDL (OxLDL) (red) were observed inside the wild-type hemocytes (left, green, *HmlΔ>GFP/+*) and *eater* mutants (right, green, *HmlΔ>GFP/+, eater*^1^) **(f)**. Quantification of the bound oxidized LDL in the hemocytes. Bounded amount was normalized to *Canton S* hemocytes **(g)**. Mann- Whitney test was performed for data analysis.

**h.** Quantification of total sleep time in the *crq* mutant (*Crq∆*) fly (left; male, right;female).

Mann-Whitney test was performed for data analysis.

**i-j.** Immunoprecipitation of acetylated GLaz from wild-type or *eater* mutant brain. GLaz expression level in the wild-type (*CantonS*, C.S) or *eater* mutant (*eater*^1^) is similar in the lysate. The level of acetylated-GLaz is also similar **(i)**. This is supported by quantification of acetylated-GLaz levels **(j)**. IB: immunoblot. IP: Immunoprecipitation. Lys^AC^: acetylated lysine. Tub: alpha tubulin. Mann-Whitney test was performed for data analysis.

**k.** Measurement of NADH levels in *eater* mutant (*eater*^1^) and wild-type (*CantonS*) brains.

Unpaired t test was performed for data analysis.

**l.** Graphs comparing total sleep time in female wild-type (*Repo-GeneSwitch(G.S)/+*), sirtuin overexpression (*Repo-GeneSwitch(G.S) UAS-Sirt1/+*), and *eater* mutant with sirtuin overexpression (*Repo-GeneSwitch(G.S) UAS-Sirt1/+, eater*^1^) in glial cells. The red shade indicates the sleep data for flies fed with RU486. Tukey’s multiple comparisons test was performed for data analysis.

**m.** Visualization of ROS using fluorescence probes in the brain. ROS dyes (Left, MitoSox, Right, DHE) were co-localized with cortex glial cell marker (green, *NP2222-Gal4 UAS- mCD8GFP/+)*.

**n.** Staining of cleaved Dcp1 as a cell-death marker in the fly brain. There are no differences in Dcp1 staining (green) in wild-type (*CantonS*, left) and *eater* mutants (*eater*^1^, right). Glial cell was visualized by repo antibody (magenta). ns: not significant (p>0.01); *p<0.1; **p<0.01,

***p<0.001. ****p<0.0001. Bars in graphs: the median. White scale bar, 100 μm unless otherwise indicated.

