## Supplementary Figure for "Sleep-dependent clearance of brain lipids by peripheral blood cells"

LDL in the hemocytes. Bounded amount was normalized to *CantonS* hemocytes **(g)**. Mann-Whitney test was performed for data analysis.

**h.** Quantification of total sleep time in the *crq* mutant (*CrqΔ*) fly (left; male, right;female). Mann-Whitney test was performed for data analysis.

**i-j.** Immunoprecipitation of acetylated GLaz from wild-type or *eater* mutant brain. GLaz expression level in the wild-type (*CantonS*, C.S) or *eater* mutant (*eater$^{1}$*) is similar in the lysate. The level of acetylated-GLaz is also similar **(i)**. This is supported by quantification of acetylated-GLaz levels **(j)**. IB: immunoblot. IP: Immunoprecipitation. Lys$^{AC}$: acetylated lysine. Tub: alpha tubulin. Mann-Whitney test was performed for data analysis.

**k.** Measurement of NADH levels in *eater* mutant (*eater$^{1}$*) and wild-type (*CantonS*) brains. Unpaired t test was performed for data analysis.

**l.** Graphs comparing total sleep time in female wild-type (*Repo-GeneSwitch(G.S)/+*), sirtuin overexpression (*Repo-GeneSwitch(G.S) UAS-Sirt1/+*), and *eater* mutant with sirtuin overexpression (*Repo-GeneSwitch(G.S) UAS-Sirt1/+, eater$^{1}$*) in glial cells. The red shade indicates the sleep data for flies fed with RU486. Tukey's multiple comparisons test was performed for data analysis.

**m.** Visualization of ROS using fluorescence probes in the brain. ROS dyes (left, MitoSox, right, DHE) were co-localized with cortex glial cell marker (green, *NP2222-Gal4 UAS-mCD8GFP/+).*

**n.** Staining of cleaved Dcp1 as a cell-death marker in the fly brain. There are no differences in Dcp1 staining (green) in wild-type (*CantonS*, left) and *eater* mutants (*eater$^{1}$*, right). Glial cell was visualized by repo antibody (magenta). ns: not significant (p>0.01); *p<0.1; **p<0.01, ***p<0.001. ****p<0.0001. Bars in graphs: the median. White scale bar, 100 μm unless otherwise indicated.

**Extended Data Table. 1** Table is related to statistical details in main figures and extended data figures. Each tab indicates a figure.

### Extended Data Figure 1

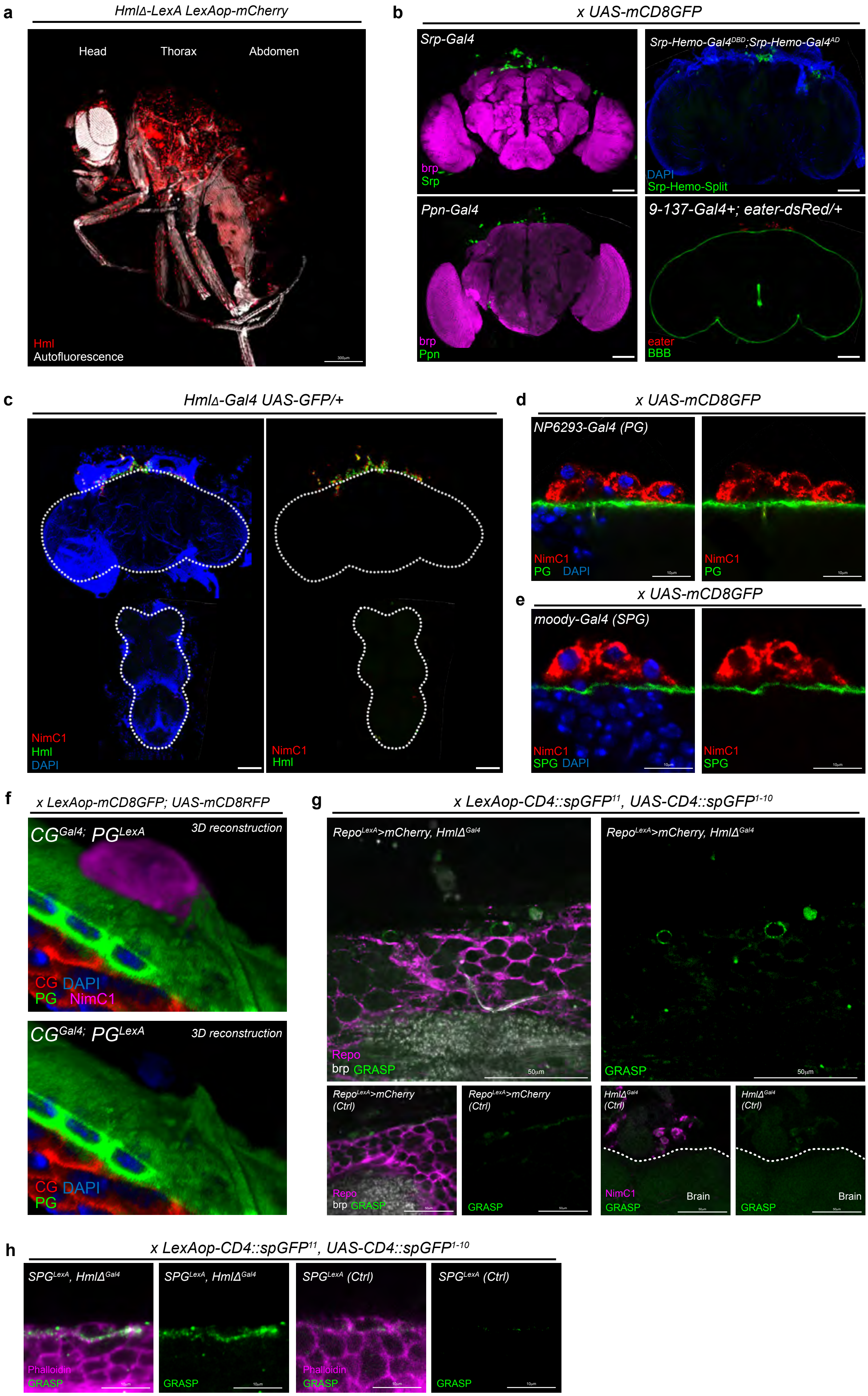

### Extended Data Figure 2

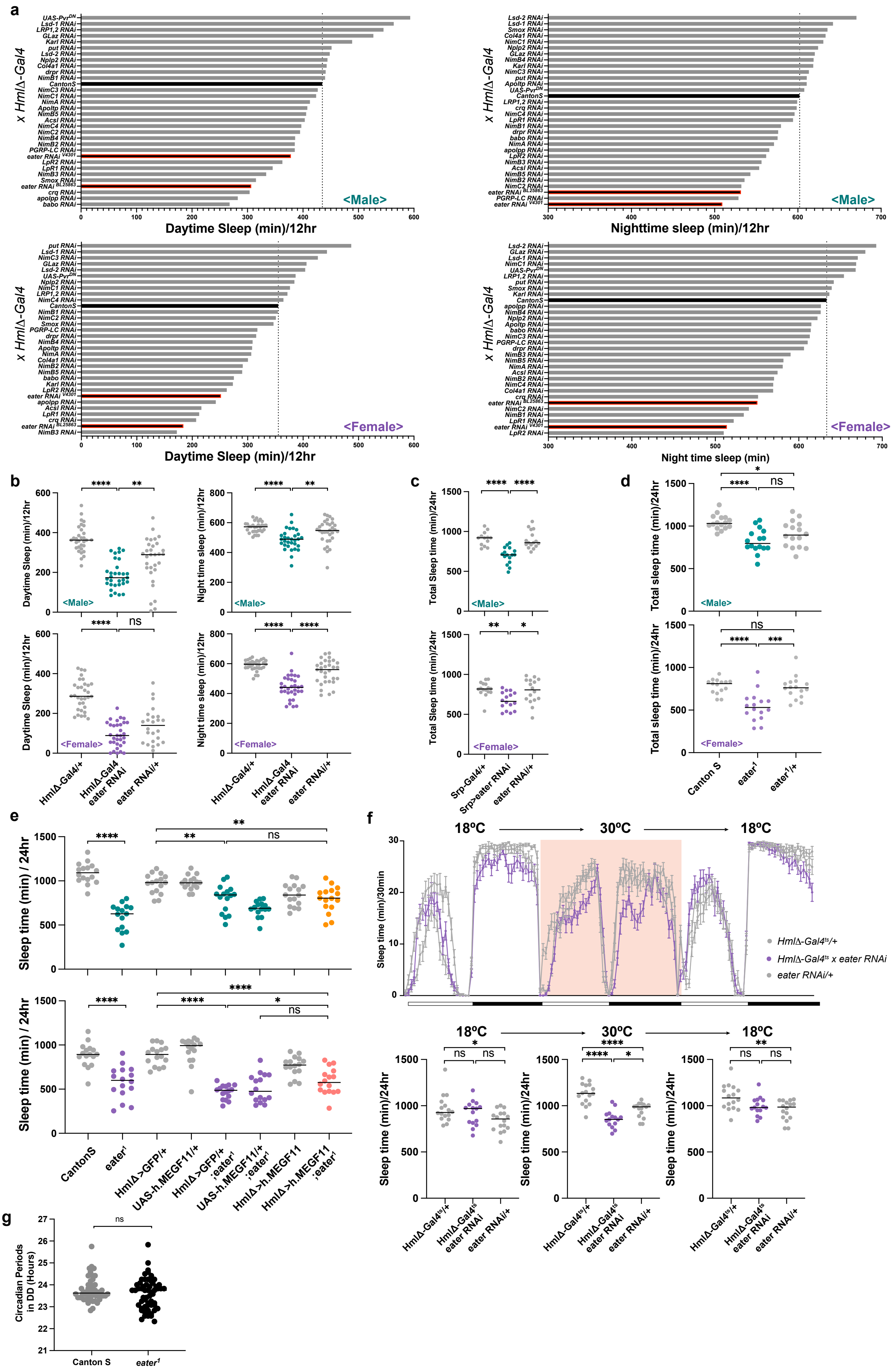

### Extended Data Figure 3

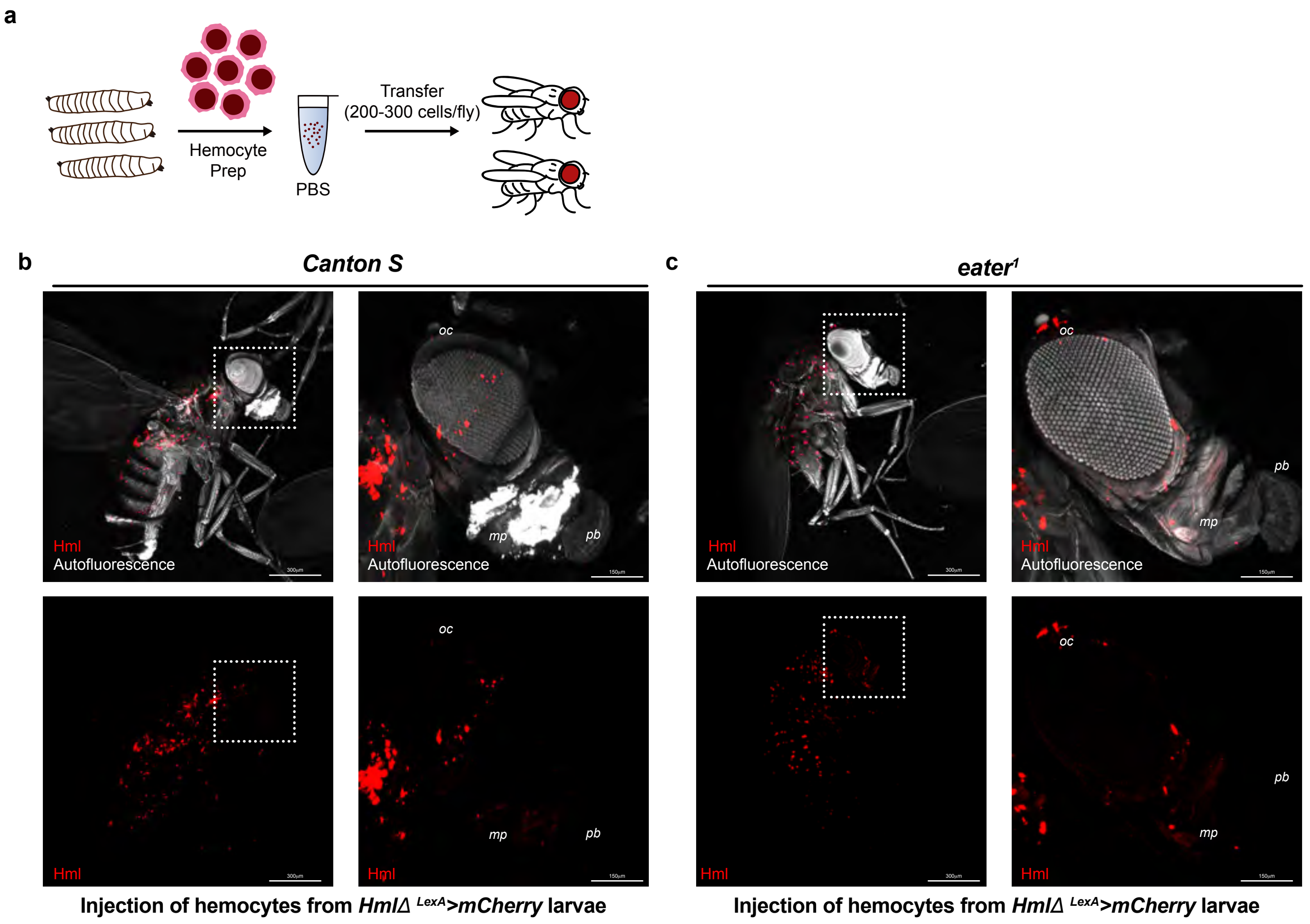

### Extended Data Figure 4

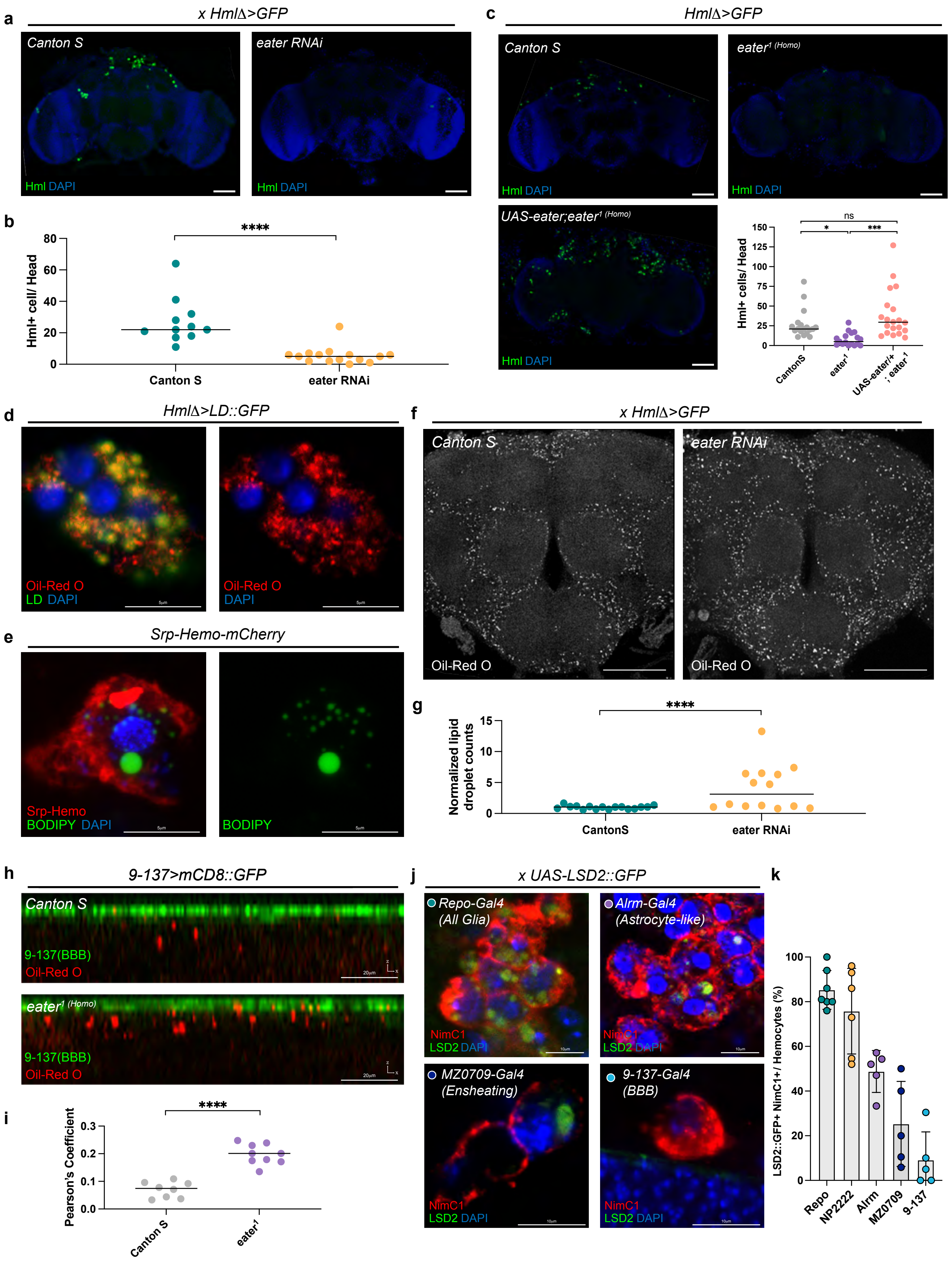

### Extended Data Figure 5

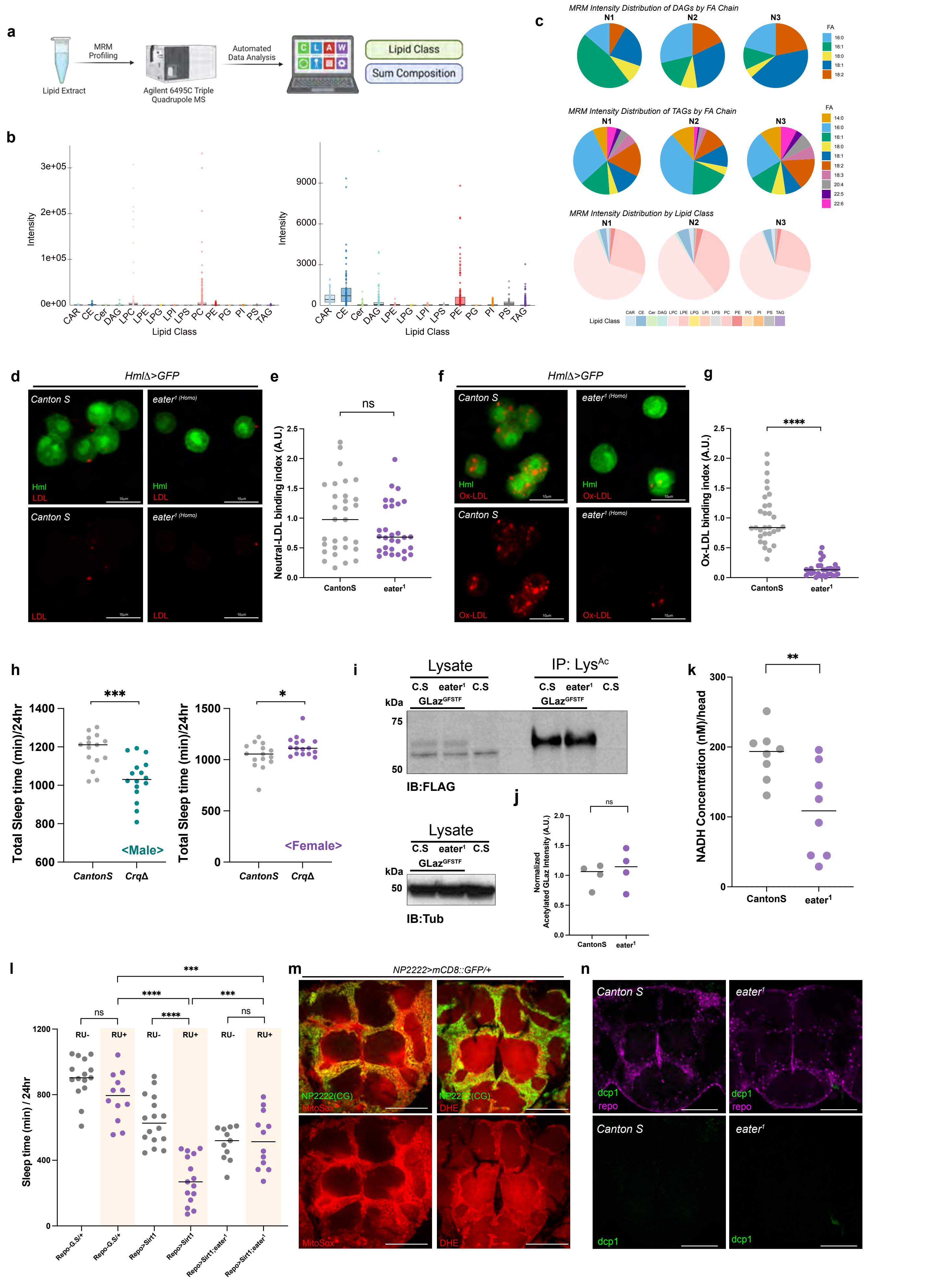
